# Short-term exposure to intermittent hypoxia leads to changes in gene expression seen in chronic pulmonary disease

**DOI:** 10.1101/2020.03.06.981241

**Authors:** Gang Wu, Yin Yeng Lee, Evelyn M. Gulla, Andrew Potter, Joseph Kitzmiller, Marc D Ruben, Nathan Salomonis, Jeffrey A. Whitsett, Lauren J Francey, John B Hogenesch, David F. Smith

## Abstract

Obstructive sleep apnea (OSA) results from episodes of airway collapse and intermittent hypoxia and is associated with a host of health complications including dementia, diabetes, heart failure, and stroke. Although the lung is the first organ to sense changes in inspired oxygen levels, little is known about the consequences of IH to the lung hypoxia-inducible factor (HIF)-responsive pathways. Furthermore, cellular mechanisms causing disease progression across multiple systems in OSA are unknown. We hypothesized that exposure to IH would lead to up- and down-regulation of diverse expression pathways and that individual cell populations would show distinctive responses to IH. We identify changes in circadian and immune pathways in lungs from mice exposed to IH. Among all cell types, endothelial cells show the most prominent transcriptional changes. Interestingly, up-regulated genes in endothelial, fibroblast, and myofibroblast cells were enriched for genes associated with pulmonary fibrosis and pulmonary hypertension. These genes include targets of several drugs currently used to treat chronic pulmonary diseases. Our results reveal potential candidates for cell-targeted therapy seeking to minimize pulmonary effects of OSA. A better understanding of the pathophysiologic mechanisms underlying diseases associated with OSA could improve our therapeutic approaches, directing therapies to the most relevant cells and molecular pathways.

## Introduction

Obstructive sleep apnea (OSA) is a condition characterized by episodes of sleep-associated upper airway obstruction and intermittent hypoxia (IH). OSA occurs in approximately 2-5% of children(Marcus et al., 2012) and 33% of adults 30-69 years of age(Benjafield et al., 2019) in the US. If untreated, OSA is associated with significant health consequences to the cardiovascular, neurological, and metabolic systems. Even young children with moderate to severe OSA can develop blood pressure dysregulation(Amin et al., 2004), systemic hypertension(Enright et al., 2003; Kohyama et al., 2003), and left ventricular hypertrophy(Amin et al., 2002, 2005). OSA is associated with a significant socioeconomic burden in the US(Sullivan, 2016). Despite available medical and surgical therapies, millions of children and adults with OSA are currently untreated or do not respond to available therapies. Cellular responses to changes in oxygen levels are primarily mediated by the hypoxia-inducible factors (HIFs). Although the lung is the first organ to sense large changes in inspired oxygen levels, little is known about the consequences of IH to the lung HIF-responsive pathways. Without a basic understanding of the molecular mechanisms that lead to diseases associated with IH and OSA, our ability to identify new treatments is significantly hindered.

Efforts to understand the effects of OSA have primarily focused on systemic inflammation(Gozal et al., 2008), oxidative stress(Tauman et al., 2014), and endothelial dysfunction(Bhattacharjee et al., 2012; Kheirandish-Gozal et al., 2013). However, the early causal events from IH-exposure are not fully elucidated. Research is now focused on other possible pathogenic pathways that could be activated or suppressed in the presence of IH, leading to disease. For example, hypoxia inducible factors (HIFs) stabilized under low oxygen conditions can affect the circadian transcriptional-translational feedback loop at the cellular level(Adamovich et al., 2017; Hogenesch et al., 1998; Kobayashi et al., 2017; Peek et al., 2017; Wu et al., 2017). Even acute exposure to IH results in dysregulation of the circadian clock that is time-of-day-dependent and tissue-specific, and these effects persist in some tissue for up to 24 hours after exposure(Manella et al., 2020). Pathways involved in immune responses and regulation can also be activated or suppressed in the presence of IH(Cubillos-Zapata et al., 2017; Lam & Ip, 2019), contributing to comorbid disease initiation and progression. Associations between IH and gene targets could be either pathogenic or protective responses for the lung. Additionally, lung could be an effector rather than target organ of IH, resulting in responses to IH that lead to multi-systemic effects.

While animal models of OSA have focused on physiologic responses to IH at organ and system levels, determination of the contributions of individual cell types in initiation and progression of disease has been challenging. Within organs, individual cells serve specific physiologic roles. As a result, pathways disrupted by stabilization of HIFs can affect cell types differently. Single-cell RNA sequencing (scRNA-seq) has emerged as a method for evaluating transcriptional states from thousands of individual cells(M. J. Zhang et al., 2018), advancing our understanding of how specific cell types contribute to physiology and disease(Plasschaert et al., 2018; M. J. Zhang et al., 2018).

In the present study, we used IH as a mouse model of OSA to better understand early cellular-specific consequences to the lung, the primary organ that first senses hypoxic episodes. We hypothesized that exposure to IH would lead to up- and down-regulation of diverse expression pathways, distinct cell populations would show distinctive responses to IH, and that changes in these gene expression pathways could provide therapeutic targets at the cell-specific level. We identify changes in both circadian and immune response pathways in lungs from mice exposed to IH. We also demonstrate strong similarities in the gene expression profiles from mice compared to those characteristics of human lung tissue from patients with diverse pulmonary diseases, including pulmonary hypertension and pulmonary fibrosis. Our results reveal potential candidates for cell-targeted therapy seeking to minimize effector responses of the lung that could lend to systemic disease. A better understanding of the pathophysiologic mechanisms underlying diseases associated with OSA could improve our therapeutic approaches.

## Results

### Short-term exposure to intermittent hypoxia reshapes circadian and immune pathways in the lung

In humans, moderate to severe OSA is associated with interstitial lung disease with remodeling of the extracellular matrix(Kim et al., 2017). Lung is the primary organ that senses episodes of hypoxia and is therefore exposed to large fluctuations in the oxygen concentrations compared to other tissues throughout the body. For these reasons, we sought to identify initial changes in gene expression pathways in the lung in response to IH.

Mice were initially entrained to the same light:dark schedule to synchronize active and inactive phases. After 14 days of entrainment in the 12h:12h light:dark cycle, mice were exposed to IH or room air (normoxia) for the entire 12h inactive phase for 9 days (Fig. 1A). Mice tolerated the procedure well. Furthermore, haemotoxylin and eosin (H&E) staining of whole lungs did not show gross changes in architecture or inflammatory remodeling after exposure to IH (Fig. S1). Bulk RNA-seq was performed to explore transcriptomic effects of IH on lung tissue at the organ level. There were 374 genes (Fig. S2A) up-regulated and 149 down-regulated in mouse lung after exposure to IH (BHQ < 0.05 and fold change >1.5). Not surprisingly, the top up-regulated genes included well-known hypoxia inducible factor (HIF)-1 target genes (e.g. *Edn1*, *Bnip3* and *Ankrd37*; Fig. S2B).

**Fig. 1.**
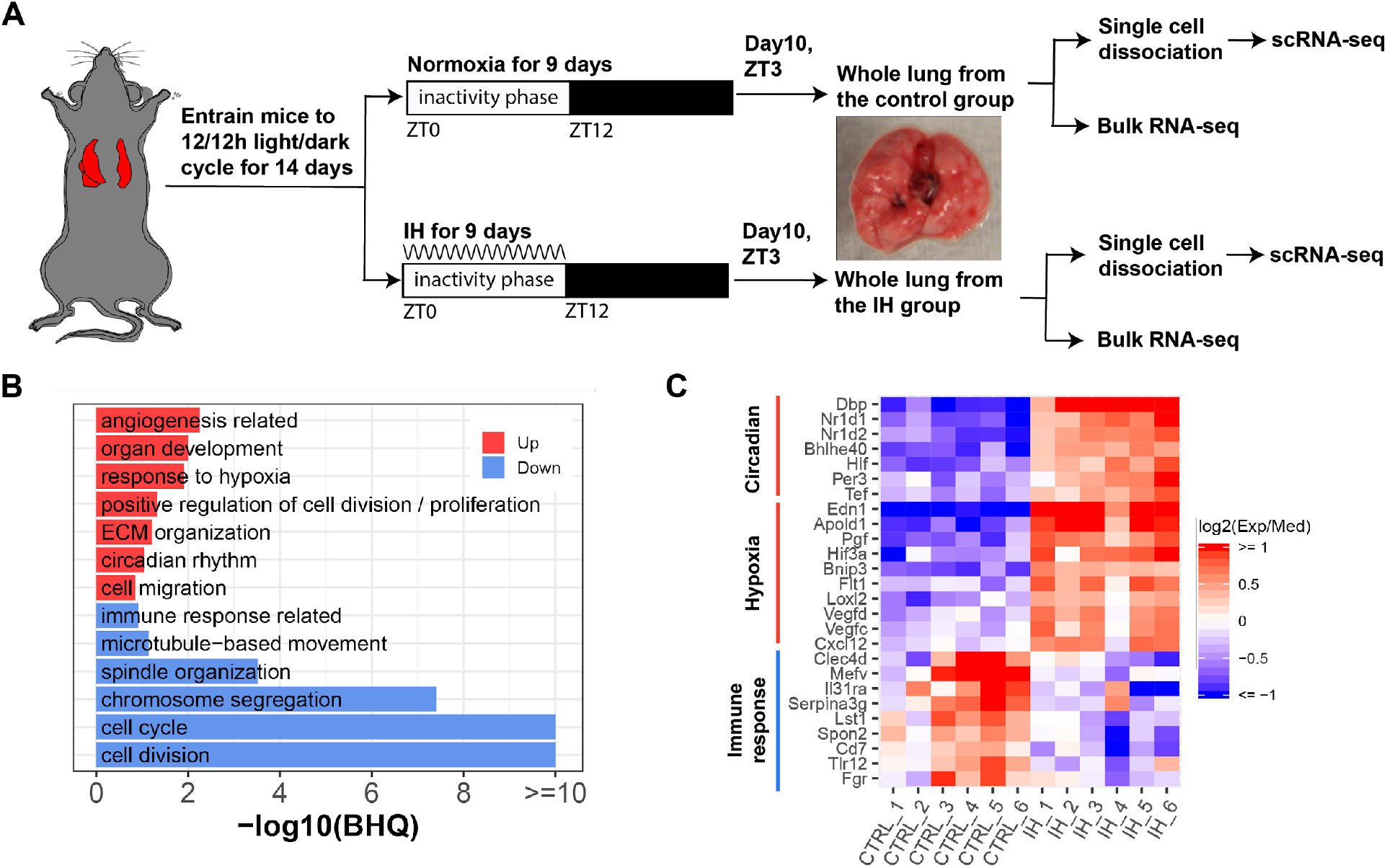
Short-term exposure to intermittent hypoxia reshapes circadian and immune pathways in the lung. (A) Schematic of IH protocol. Mice are entrained to the same 12 h:12 h light:dark cycle for 14 days prior to IH exposure. Mice are exposed to normoxia (controls) or intermittent episodes of hypoxia (21 to 6% oxygen saturation) followed by recovery to 21% oxygen over the entire 12 h inactivity phase for 9 days. Mice are then sacrificed at ZT 3 (3 h after lights-on) on the 10th day for tissue harvest. Bulk RNA-seq and scRNA-seq were performed for each group. (B) Biological processes enriched in lung from mice exposed to IH vs controls. Enrichment analysis was performed in the DAVID database, using the top 200 up and down regulated genes identified from differential expression analyses. Redundant biological processes are merged into one category. Biological processes enriched in up and down regulated genes are indicated in red and blue bars, respectively. (C) The heatmap shows the fold change of associated genes in circadian rhythm, response to hypoxia, and immune response. The red and blue indicates up and down regulated genes in the experimental group. There are six biological replicates for each group.

We performed DAVID(Huang et al., 2009) enrichment analysis to identify biological processes associated with the top 200 up and down regulated genes from mice exposed to IH. Pathways induced in response to hypoxia included circadian rhythm, angiogenesis, and extracellular matrix organization (Fig. 1B and C). Increased expression of RNAs associated with angiogenesis, such as vascular endothelial growth factor (VEGF), was observed after IH, consistent with findings in a mouse model of prolonged exposure to IH(Reinke et al., 2011) and after 72 h of IH exposure to endothelial cells *in vitro(Wohlrab et al., 2018)*. Unexpectedly, RNAs associated with immune responses were significantly down regulated after 9 days of IH (Fig. 1B and C). Present findings contrast with the general concept that HIFs are important regulators of inflammation and immune responses (Eltzschig & Carmeliet, 2011; Scholz & Taylor, 2013; C. T. Taylor et al., 2016). For example, activation of neutrophils by HIFs is largely considered proinflammatory(Peyssonnaux et al., 2005; C. T. Taylor et al., 2016; Walmsley et al., 2005). As previously reported, there is a tight interaction between clock genes and HIFs(Edgar et al., 2012; Gu et al., 2000; Hogenesch et al., 1998; McIntosh et al., 2010; B. L. Taylor & Zhulin, 1999). Several circadian clock repressors (e.g. *Nr1d1*, *Nr1d2*, *Bhlhe40* and *Per3*) were significantly upregulated in the IH group (Fig. 1C).

### Single cell sequencing identifies 19 distinct cell types in the lungs of intermittent hypoxia and control mice

We detected significant differences related to a number of cell-type selective genes on biological functions. We then applied single cell transcriptomics to identify cell-type specific effects of IH. In total, we sequenced 12,324 and 16,125 pulmonary cells from IH and control mice. Unsupervised analysis identified 25 transcriptionally distinct cell clusters, corresponding to 19 distinct cell types (Fig. 2A-C) based on the expression of established marker genes (See Methods), including stromal, epithelial, endothelial, immune, as well as small numbers of other cell types. The proportion of endothelial, AT2, fibroblast/myofibroblast cells were modestly increased, but the proportion of immune cells (e.g. B and T cells) was decreased in the IH exposed mice (Fig. 2C). Overall, the variation of lung cell types was small (BHQ > 0.05). Given that this was a short exposure of IH, we did not anticipate a dramatic difference in cell types or proportions. We then examined samples to determine if lungs from control vs experimental mice showed histologic differences in cell types.

**Fig. 2.**
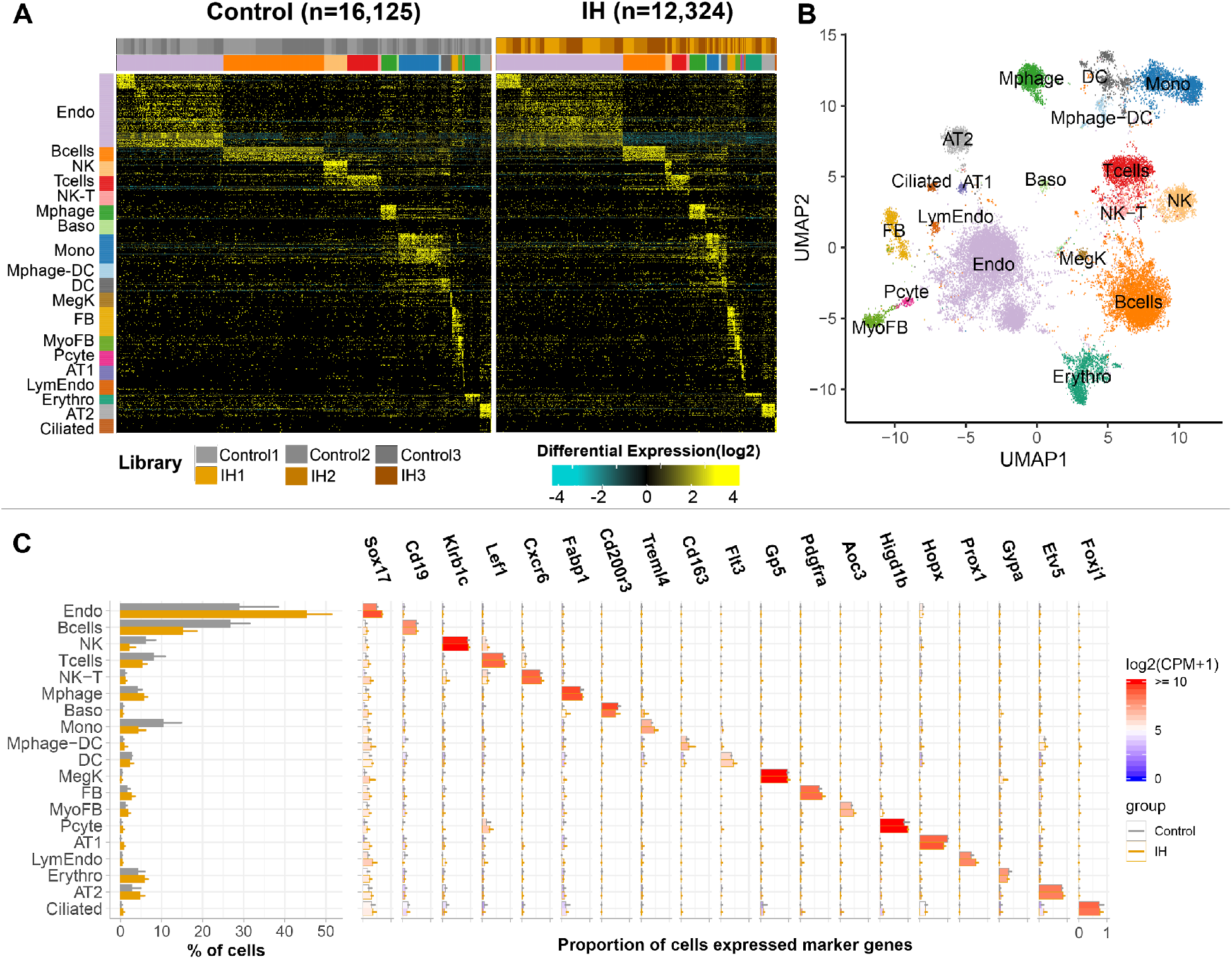
Single cell sequencing identifies 19 distinct cell types in the lungs of intermittent hypoxia and control mice. (A) Heat map of AltAnalyze selected marker genes for each cell population. Columns and rows represent individual cells and marker genes, respectively. Biological replicates and cell types are indicated by the color bars on the top. cellHarmony was used to align cells from control to IH groups. (B) UMAP projection of whole lung cell populations from IH and control mice in the heatmap. (C) Percentage (left panel) and expression of marker genes (right panel) for each cell type from experimental and control mice. Error bars indicate standard deviation of cell proportion from the three replicates. The list of cell types include: endothelial cells (Endo), B cells (Bcells), natural killer cells (NK), T cells (Tcells), natural killer cells (NK), natural killer T cells (NK-T), macrophages (Mphage), basophils (Baso), monocytes (Mono), macrophages-dendritic CD163+ cells (Mphage-DC), dendritic cells (DC), megakaryocytes (MegK), fibroblasts (FB), myofibroblasts (MyoFB), pericytes (Pcyte), alveolar type I cells (AT1), lymphatic endothelial cells (LympEndo), erythroblasts (Erythro), alveolar type II cells (AT2), and ciliated cells (Ciliated). CPM indicates UMI count per million.

### Short-term exposure to intermittent hypoxia did not lead to histologic changes in the lung

Similar to H&E histology, confocal immunofluorescence microscopy for major cell types were normal without inflammatory remodeling. Immunostaining for endothelial markers LYVE1 and FOXF1 did not show changes in IH exposed mice (Fig. 3C-D) compared to normoxia (Fig. 3A-B). Also important, staining for MKI67 did not show changes in endothelial cell proliferation (Fig. 3E-H). Expression levels of HOPX and SFTPC in alveolar type I and type II cells were not significantly different for IH (Fig. 3K-L) versus control (Fig. 3I-J) mice. Immunostaining for the progenitor marker SOX9 (Fig. 3I-L) or the extracellular matrix marker POSTN (Fig. 3M-P) did not demonstrate any changes after IH exposure. Overall, there were no differences in endothelial or epithelial cells, signs of fibrosis, or increases in the number of progenitor or proliferating cells. Although other studies have demonstrated changes in proliferating type II alveolocytes(Reinke et al., 2011) and pulmonary vascular remodeling(Nisbet et al., 2009), these were chronic mouse models of IH involving months of exposure. These models also produced other phenotypic changes, such as increased lung volumes(Reinke et al., 2011) and pulmonary hypertension(Nisbet et al., 2009). We exposed our mice to IH for a shorter period of time to specifically evaluate the changes in gene expression prior to lung remodeling with the hope of uncovering early pathways that lead to disease.

**Fig. 3.**
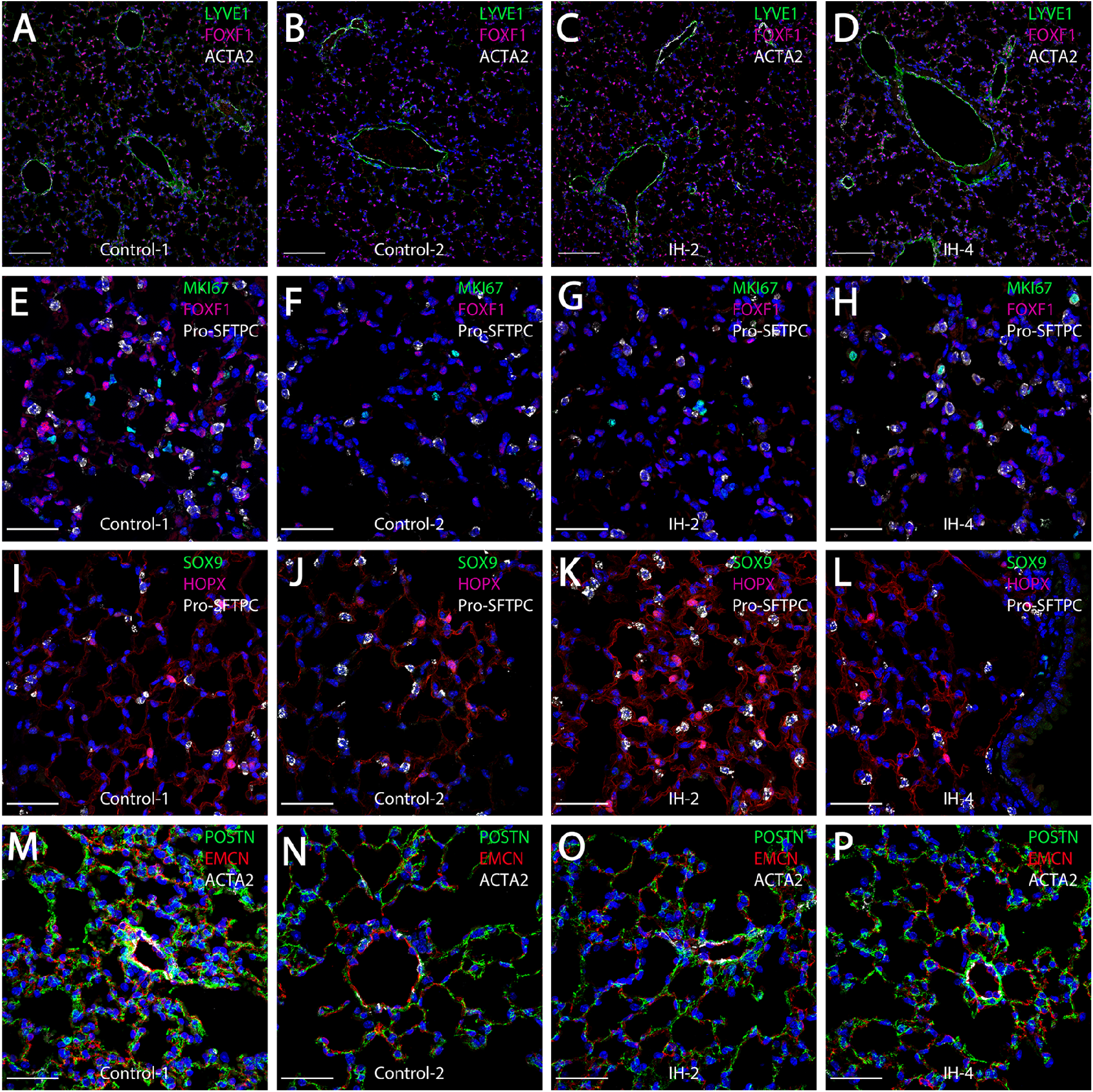
Short-term exposure to intermittent hypoxia did not lead to histologic changes in the lung. (A-D) Immunostaining for LYVE1, FOXF1, and ACTA2 in mouse lung show no changes in the number or clustering of endothelial cells. (E-H) Immunostaining for MKI67, FOXF1, and Pro-SFTPC did not show changes in the number of proliferating endothelial cells. (I-L) Expression levels of SOX9, HOPX, and Pro-SFTPC did not demonstrate differences in the number of progenitor cells, alveolar type I cells, or alveolar type II cells. (M-P) Immunostaining for POSTN, EMCN, and ACTA2 in mouse lung did not show IH-induced changes in extracellular matrix markers. The scale bars for A-D represent 100 μm. The scale bars for E-P represent 40 μm. KEY: LYVE1 is expressed at high levels in lymphatic and vascular endothelial cells. FOXF1 and ACTA2 are expressed at high levels in vascular endothelial and smooth muscle cells, respectively. MKI67 is a marker for cell proliferation. HOPX and Pro-SFTPC show high expression levels in the alveolar type I cells and alveolar type II cells, respectively. SOX9 is a marker for progenitor cells. POSTN is used to stain extracellular matrix. EMCN is a marker for capillaries and veins/venules.

### Diverse expression pathways were up and down regulated in the presence of intermittent hypoxia

We further explored the early cell-type specific response to IH in mouse lung by aggregating single cell data into “pseudo-bulk” data to compare biological replicates for each identified cell type (see Methods for details). Using DESeq2(Love et al., 2014), the number of up or down regulated genes in different lung cell types in response to IH are not equal (Fig. S3). To balance the difference in the pathway enrichment analysis, we selected the top 200 up and down regulated genes (ranking by the *P* value) in each cell type. From the DAVID enrichment analysis, diverse biological processes were up and down regulated in different cell types in response to IH (Fig. 4A, Fig. S4 A and B). For example, hypoxia-responsive and circadian pathways were enriched in those up regulated genes in response to IH in endothelial cells, myofibroblasts, and AT2 cells. Immune response-related and antigen processing and presentation were enriched in those down regulated genes in monocytes, macrophages-dendritic cells, NK cells, and erythroblasts. Surprisingly, circadian pathways were highly enriched in multiple cell populations, not just epithelial cells, a population that is important for circadian rhythmicity in the lung(Gibbs et al., 2009). As expected from specific genes in each biological process, the response level of the same genes were different in multiple cell types (Fig. 4B and C). For example, the circadian repressor gene, *Nr1d1,* was more responsive to IH in endothelial and AT2 cells than lymphatic endothelial cells and AT1 cells. We also noted cell-specific responses for the down regulated genes. For example, immune response genes (e.g. *Iglc3*, *S100a8*, and *Oas3*) decreased more in monocytes than macrophages in response to IH.

**Fig. 4.**
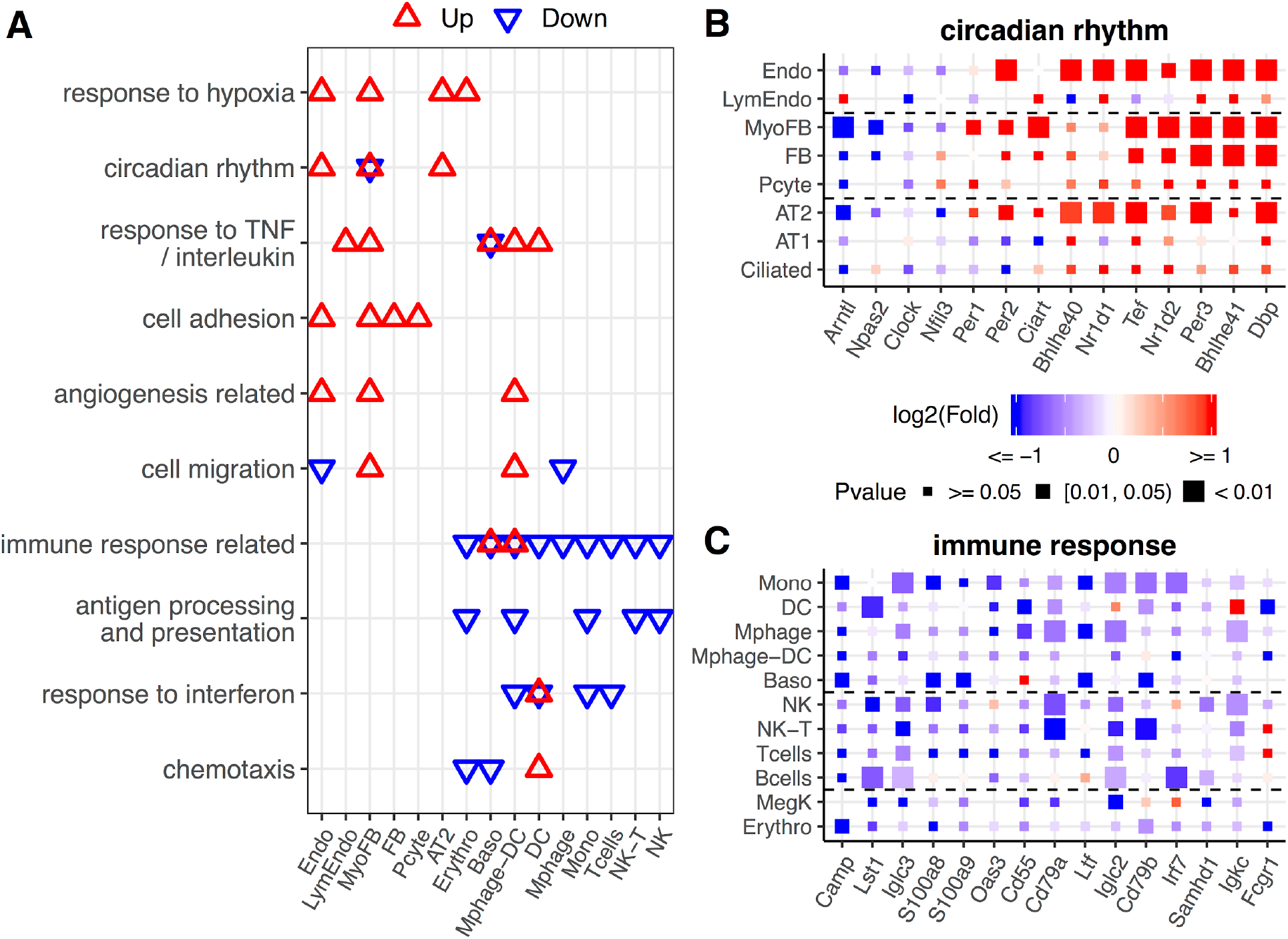
Diverse expression pathways were up and down regulated in the presence of intermittent hypoxia. (A) Biological processes enriched in different cell types from lungs of mice exposed to IH vs Control. Enrichment analysis was performed in the DAVID database, using the top 200 up and down regulated genes identified from each cell type. Redundant biological processes are merged into one category. Biological processes enriched in up and down regulated genes are indicated in red and blue triangles, respectively. (B) Expression variation of well-established genes involved in circadian rhythm for endothelial, epithelial, and mesenchymal cells. (C) Expression variation of well-established genes involved in immune response for immune associated cells. For B and C, the fold change is indicated by the color, and the P-value for differential expression is indicated by the point size. The list of cell types include: endothelial cells (Endo), B cells (Bcells), natural killer cells (NK), T cells (Tcells), natural killer cells (NK), natural killer T cells (NK-T), macrophages (Mphage), basophils (Baso), monocytes (Mono), macrophages-dendritic CD163+ cells (Mphage-DC), dendritic cells (DC), megakaryocytes (MegK), fibroblasts (FB), myofibroblasts (MyoFB), pericytes (Pcyte), alveolar type I cells (AT1), lymphatic endothelial cells (LympEndo), erythroblasts (Erythro), alveolar type II cells (AT2), and ciliated cells (Ciliated).

### Pulmonary vascular endothelial subpopulations show distinctive responses to intermittent hypoxia

Recent studies show distinct vascular endothelial cell subpopulations in mouse and human lung(Ren et al., 2019). Our vascular endothelial populations were annotated to endothelial artery, vein, capillary aerocytes (Cap-a), and general capillary (Cap-g) cells (Fig. 5A and Fig. S5). Interestingly, we found that endothelial cells demonstrated profound changes in gene expression profiles in response to IH. The endothelial capillary cells were more responsive to IH compared to endothelial artery and vein cells (Fig. 5B). For example, at BHQ < 0.2, more than 100 genes were significantly up-regulated in capillary aerocytes and general capillary cells. However, only 1 gene in the arterial endothelial cells and 57 genes in the venular endothelial cells were significantly up-regulated at the same cutoff. This trend persisted at other BHQ cut-offs (Fig. S6). General capillary cells are more responsive to hypoxia than capillary aerocytes. In addition, more genes were significantly up and down regulated in general capillary cells (Fig. 5B). Hypoxia-responsive genes (e.g. *Pdk1*, *Vegfc*, *Slc2a1*, *Pkm*, and *Flt1*) showed higher levels of expression variation (fold change and significance) in general capillary cells than capillary aerocytes (Fig. 5C), demonstrating variation at the subpopulation level. Glycolytic process was up regulated and cell migration was down regulated in both capillary aerocytes and general capillary cells in response to IH (Fig. 5D and E). Given the association between glycolysis, cytoskeletal remodeling, and cell migration in other cell types(Shiraishi et al., 2015), the similar enrichment trends for these pathways is not surprising. Additionally, without vascular growth associated with chronic IH, glycolysis may be used to meet metabolic demand. Interestingly, changes in the glycolytic process were specific to endothelial cell types (Fig. S4B).

**Fig. 5.**
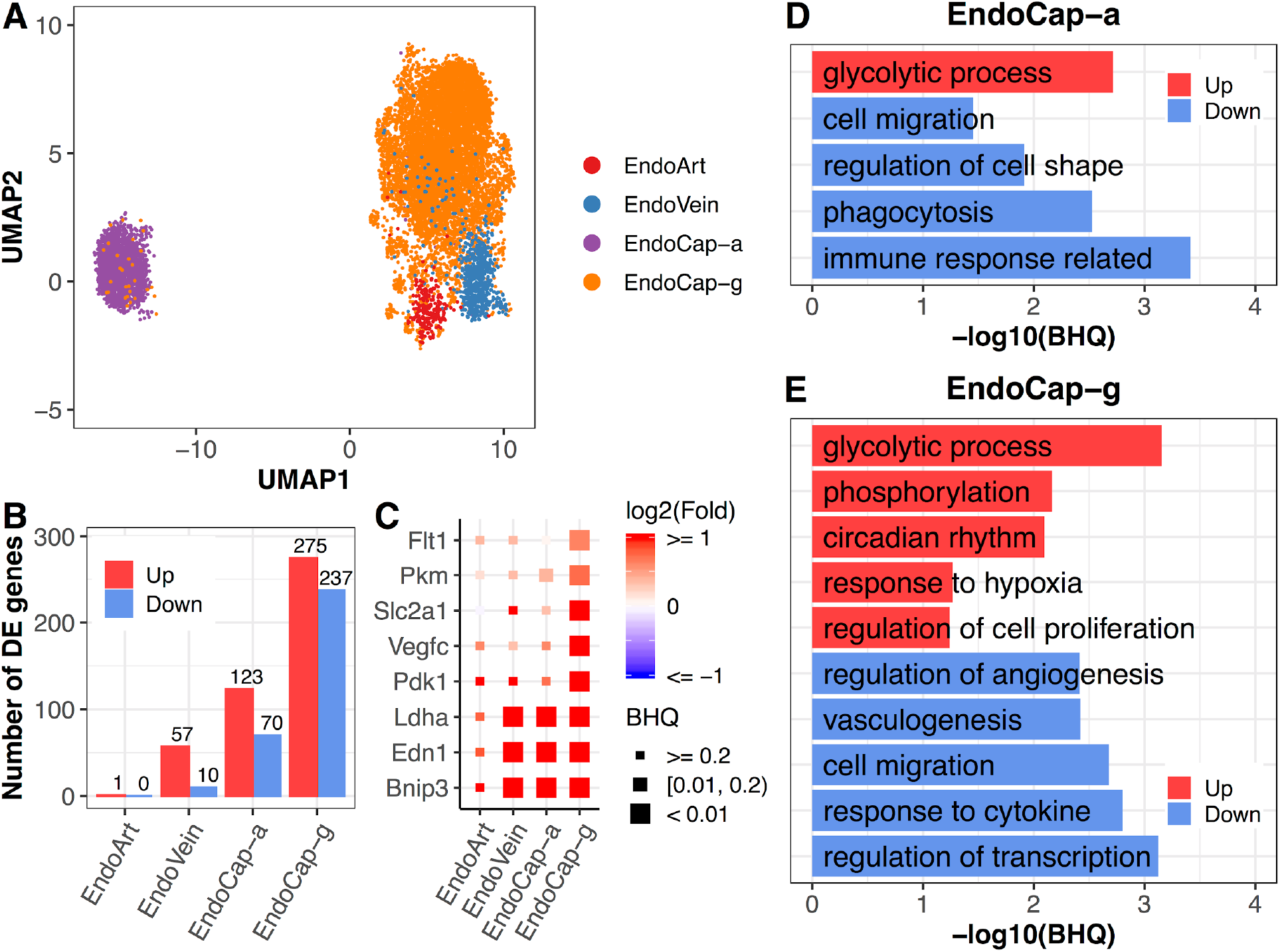
Pulmonary vascular endothelial subpopulations show distinctive responses to intermittent hypoxia. (A) UMAP projection of cells from four lung endothelial subpopulations. (B) The number of differentially expressed (BHQ < 0.2) genes in each endothelial subpopulation from IH vs control. (C) Expression variation of related genes in response to IH is shown for each endothelial subpopulation. Biological processes enriched in capillary aerocyte (D) and capillary general (E) from IH vs control mice. Enrichment analysis was performed in the DAVID database, using up and down regulated genes (BHQ < 0.2). Redundant biological processes are merged into one category. Top 5 biological processes enriched in up and down regulated genes are indicated in red and blue bars, respectively. The list of endothelial subpopulations include endothelial artery (EndoArt), vein (EndoVein), capillary aerocytes (EndoCap-a), capillary general (EndoCap-g) cells.

On the other hand, capillary aerocytes and general capillary cells demonstrated more differences in enrichment pathways (Fig. 5D and E). For example, circadian rhythm and regulation of angiogenesis were only enriched for those up and down regulated genes in general capillary cells in response to IH, while phagocytosis and regulation of cell shape were only enriched for those down regulated genes in capillary aerocytes (Fig. 5D and E, table S12). Anatomic location within the pulmonary vasculature, including variable roles in gas exchange, could lead to these differences in response. Alternatively, proximity and interaction with other cell types, such as fibroblasts or immune cells, could also help to explain these findings.

Evaluation of the expression pathways at the single cell resolution demonstrated significant changes in multiple cell types. With this information, we wanted to identify potential candidates for therapeutic intervention in response to IH.

### Pulmonary disease-regulated genes provide clinical implications for OSA at the cell-specific level

OSA is associated with an array of pulmonary diseases, such as interstitial lung disease (Kim et al., 2017), idiopathic pulmonary fibrosis (Lancaster et al., 2009), and pulmonary hypertension (Chaouat et al., 1996; Dimitar Sajkov & McEvoy, 2009). IH led to significantly more up than down regulated genes (Fig. 6A and Fig. S7). IH-induced expression in myofibroblasts, fibroblasts, AT2 cells, basophils, and macrophage-DCs were significantly enriched for chronic obstructive pulmonary disease (COPD)-associated genes, while myofibroblasts and fibroblasts demonstrated enrichment of genes associated with pulmonary fibrosis. However, the disease genes were not equally expressed or up regulated in these cell types (Fig. 6B). For example, *Ptgis* is a pulmonary hypertension-associated gene highly expressed in myofibroblasts and fibroblasts compared to other cell types (Fig. S8). *Ptgis* is also a target gene for Epoprostenol–– a drug used for treating pulmonary hypertension(Sitbon & Vonk Noordegraaf, 2017). The idiopathic pulmonary fibrosis-associated gene, *Thbs1(Idell et al., 1989; Kuhn & Mason, 1995; Yehualaeshet et al., 2000)*, was highly expressed and more responsive to IH in myofibroblasts compared to other cell types. *Msr1(Hersh et al., 2006; Silverman et al., 2002)* is a COPD-associated gene which was highly expressed in macrophage-DCs, basophils, and monocytes. These data highlight the similarity of the IH signatures with cell-type specific responses in an array of pulmonary diseases (Fig. 6C, Fig. S8).

**Fig. 6.**
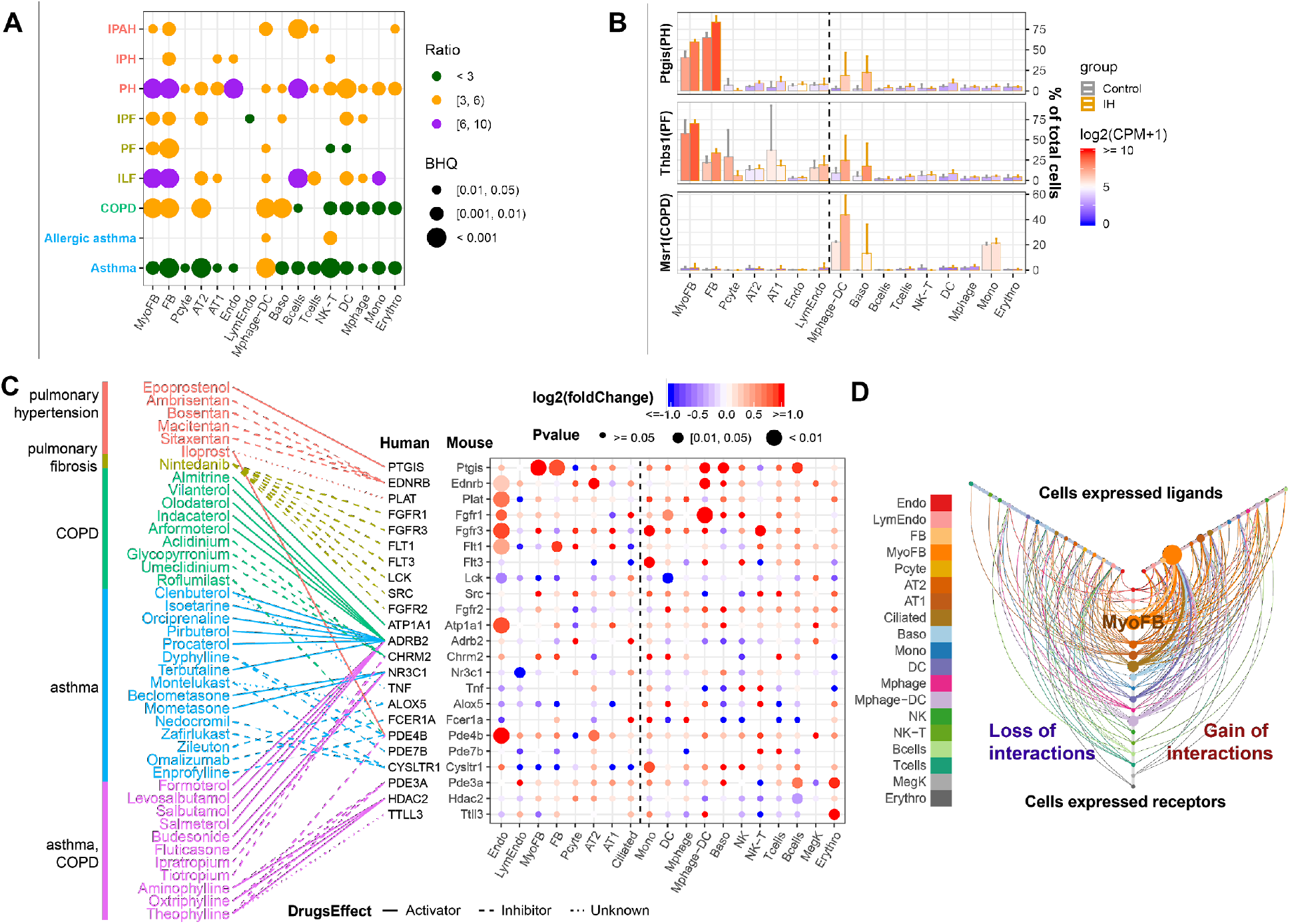
Pulmonary disease-regulated genes provide clinical implications for OSA at the cell-specific level. (A) Pulmonary disease-associated genes are enriched in up regulated genes in different cell types from whole lungs of mice exposed to IH vs controls. Enrichment analysis was performed in DisGenet human database, using the top 200 up regulated genes identified from each cell type. The enrichment ratio is indicated by color. The point size indicates the enrichment BHQ from a fisher exact test. The pulmonary diseases include asthma, allergic asthma, chronic obstructive airway disease (COPD), interstitial lung fibrosis (ILF), pulmonary fibrosis (PF), idiopathic pulmonary fibrosis (IPF), pulmonary hypertension (PH), idiopathic pulmonary hypertension (IPH), and idiopathic pulmonary arterial hypertension (IPAH). (B) The disease-associated genes vary in expression level and percentage of cells that express those genes. The fold change is indicated by the color, and the percentage of cells that express those genes is indicated by the height of the bar. Control and experimental groups are indicated by grey and orange box borders, respectively. (C) Dozens of pulmonary drug targets show differential expression in multiple cell types in lung from mice exposed to IH. Drug classes used to treat different pulmonary diseases are indicated by text color. Drug effect is indicated by the line type. Fold change is indicated by the point color, and the P-value of differential expression is indicated by the point size. (D) The hive plot shows ligand-receptor interaction changes between pairs of cell types in response to IH. Nodes indicate cell-expressed ligands (horizontal axis) or receptors (vertical axis). Size of nodes are in proportion to the number of interactions changed for the cell type. Width of lines show numbers of interactions gained (right) or lost (left) between the pairs of cell types. The list of cell types include: endothelial cells (Endo), B cells (Bcells), natural killer cells (NK), T cells (Tcells), natural killer cells (NK), natural killer T cells (NK-T), macrophages (Mphage), basophils (Baso), monocytes (Mono), macrophages-dendritic CD163+ cells (Mphage-DC), dendritic cells (DC), megakaryocytes (MegK), fibroblasts (FB), myofibroblasts (MyoFB), pericytes (Pcyte), alveolar type I cells (AT1), lymphatic endothelial cells (LympEndo), erythroblasts (Erythro), alveolar type II cells (AT2), and ciliated cells (Ciliated).

We further evaluated changes in the interactions of these drug-targeted disease genes in each cell type using CellPhoneDB(Vento-Tormo et al., 2018). There are 32 ligand-receptor interactions found in the database using these drug-targeted disease genes. Overall, myofibroblasts were involved in 98 out of 176 gains of interaction (BHQ=4.28e-50), demonstrating the significance of this cell type in activating the FGF signaling pathway in early IH-associated responses (Fig. 6D, Fig. S9).

## Discussion

OSA results from intermittent episodes of airway collapse and hypoxemia and is associated with dementia(Osorio et al., 2015), diabetes(Punjabi & Beamer, 2009), hypertension(Marin et al., 2012), heart failure(Gottlieb et al., 2010), and stroke(Valham et al., 2008). How cellular responses to hypoxia and hypoxemia initiate and cause disease progression in multiple organs remains unknown. Using IH as a mouse model of OSA, we show profound, cell-type specific changes in genome-wide RNA expression in the lung. RNA profiles from lungs of mice exposed to IH shared similarity with gene expression changes in human lung from patients with pulmonary disease, including pulmonary hypertension, COPD, and asthma. OSA is associated with injury to alveolar epithelial cells and extracellular matrix remodeling, key features of interstitial lung disease (Kim et al., 2017). Although it is known that pulmonary diseases share general mechanisms, such as systemic inflammation and oxidative stress(McNicholas, 2009), there is an incomplete understanding of the early-stage changes in the lung from OSA.

In the present study, macrophages, dendritic cells, and NK cells were the only populations to demonstrate altered oxidation-reduction in the early stages of IH exposure. It was previously demonstrated that chronic IH in mice caused the release of free oxygen radicals in the lung(Tuleta et al., 2016), effects that could be blunted with antioxidative agents(Tuleta et al., 2016). Pulmonary hypertension from chronic IH was associated with enhanced NADPH oxidase expression, and knockout mice lacking one of these subunits demonstrated attenuated effects of chronic IH(Nisbet et al., 2009). Which cells mediate these effects in the lung? This type of knowledge can help direct therapeutics to the most relevant cells and molecular pathways.

In our mice exposed to IH, prostacyclin synthase (*Ptgis*) expression was dramatically upregulated in myofibroblasts and fibroblasts, and to a lesser degree macrophages. As a potent vasodilator and inhibitor of platelet aggregation, therapies targeting this pathway are already used for patients with pulmonary arterial hypertension(Farber & Gin-Sing, 2016). Given the association between OSA and pulmonary hypertension(Bady et al., 2000; D. Sajkov et al., 1999), further work can determine the clinical potential from targeting these pathways. As another example, IH downregulated fibroblast growth factor receptor (FGFR-2) in alveolar type 2 cells. FGFR2 promotes alveolar regeneration in response to lung injury(Perl & Gale, 2009), and is upregulated in patients with idiopathic pulmonary fibrosis(Li et al., 2018). Previous studies suggest that OSA can induce injury to the lung(Aihara et al., 2011; Lederer et al., 2009), and OSA is prevalent in patients with idiopathic pulmonary fibrosis(Lancaster et al., 2009). If OSA does, in fact, lead to fibrotic changes in the lung, targeting FGF pathways in alveolar epithelial cells could prevent disease progression from IH.

Although there is limited data on the role of the circadian clock in OSA(von Allmen et al., 2018), it may play a significant role in this disease(Butler et al., 2015; Entzian et al., 1996; Smith et al., 2017). We found dysregulation of circadian gene expression in multiple cell types. The circadian clock is a transcriptional/translational feedback loop that coordinates 24 h timing of physiological functions. BMAL1, the key clock transcription factor(Hogenesch et al., 1998), interacts with CLOCK(King et al., 1997) and it’s partner NPAS2(Zhou et al., 1997) to activate hundreds of target genes. All three are members of the basic helix-loop-helix (bHLH)-PER-ARNT-SIM (PAS) transcription factor family. The HIFs (1-3) are members of the same transcription factor family and are stabilized under low oxygen conditions. It is known that the circadian clock in alveolar epithelial cells impacts pulmonary physiology, and it’s disruption can contribute to disease in animal models(Z. Zhang et al., 2019). More importantly, IH leads to intertissue circadian misalignment (including in the lung) in mice(Manella et al., 2020). The hypoxic episodes that define OSA are clearly diurnal, but we don’t understand if clock disruption is a cause or consequence of disease. Our findings suggest that circadian clock dysfunction may be an important early-stage consequence of hypoxia-driven disease and may contribute to downstream processes.

Lung samples from IH-exposed mice did *not* show comprehensive histopathologic changes. This contrasts with findings from some prior murine models of IH. For example, IH induced epithelial cell proliferation in lungs from a chronic exposure model (Reinke et al., 2011). In a bleomycin-induced lung injury model, fibrosis in mouse lung is worsened by IH(Gille et al., 2018). In our model, short-term exposure to IH did not result in changes to the parenchyma or vessels. This is likely attributed to the difference in the length of time of IH exposure. Future longitudinal studies should address how gene expression profiles change over time and which cell types drive disease progression from early to late IH exposure. Early insults from hypoxia may also drive organ-specific damage in other systems. Although models of IH replicate desaturation and recovery of fractional inhaled oxygen, there is variability in the number of hypoxic events, length of desaturation events, and length of overall exposure. This variability could affect expression profiles and histopathologic findings.

Our data provide insight into the early cellular responses that drive disease progression in OSA. By identifying the roles of individual cells in disease, we have the opportunity to test targeted therapeutics, focusing specifically on the most pathologically-relevant cells and molecular pathways. Studies using scRNA-seq are already being used to identify novel cell populations in disease. For example, Xu and colleagues(Xu et al., 2016) identified a loss of normal epithelial cells in the development of idiopathic pulmonary fibrosis. In another study using single-cell profiling of bronchial epithelial cells, the major source of cystic fibrosis transmembrane conductance regulator (CFTR) activity, the pulmonary ionocyte, was revealed(Plasschaert et al., 2018). As *CFTR* is the gene mutated in cystic fibrosis, cell-specific therapies for this disease can now be evaluated.

Delineating the upstream processes dysregulated in OSA could help us identify potential candidates for therapeutic intervention. Given the socioeconomic burden to our healthcare system for diagnosing and treating OSA, new diagnostic and therapeutic strategies will be vital for the coming years.

## Materials and Methods

### Animals

Use of animals and all procedures were approved by the Institutional Animal Care and Use Committee at Cincinnati Children’s Hospital Medical Center and complied with the National Institutes of Health guidelines. Male C57BL/6J wild type mice aged 6 weeks were purchased from The Jackson Laboratory (Bar Harbor, ME) and entrained to a 12h:12h light dark cycle for 2 weeks prior to exposure.

### Experimental Design

All studies were conducted in 8-10 week old male mice. Mice were housed in light boxes and entrained to a 12:12 light:dark cycle for 2 weeks prior to initiation of IH. Mice were randomly assigned to IH or room air exposures. For the experimental group, mice were maintained in a commercially-designed gas control delivery system (Model A84XOV, BioSpherix, Parish, NY) during the inactive (light) phase from (ZT 0-12). Mice were provided with food and water *ad libitum*. For each episode of IH, the fractional inhaled oxygen (O_2_) was reduced from 20.9% to 6% over a 50 s exposure period, followed by an immediate 50 s recovery period to 20.9%. The fractional oxygen was maintained at 20.9% for approximately 15 s before the cycle was repeated, allowing for ~30 hypoxic events per hour. Ambient temperature in the hypoxia chamber was maintained between 22-24°C to match room air. Mice in the experimental group were maintained at room air during the active phase, ZT 13-24. Mice in the control group were maintained at room air throughout the circadian cycle, ZT 0-24. Experimental and control mice were exposed to IH vs room air for 9 days, followed by sacrifice at ZT 3 on day 10 of exposure. This was immediately followed by organ harvest and preparation for bulk RNA-seq, scRNA-seq, or histopathology.

### RNA Isolation

In total, 6 experimental and 6 control mice were used for bulk RNA-seq. For bulk RNA-seq, the lung was quickly harvested and snap-frozen in liquid nitrogen. Organs were later homogenized in TRIzol reagent (Invitrogen) and processed using a bead mill homogenizer (Qiagen Tissuelyser). RNA was then isolated from lung homogenates by phase separation using chloroform and phase separation columns. The aqueous phase was then applied to an RNeasy column following the manufacturer’s protocol (Qiagen) to extract and purify the RNA.

### Bulk RNA Sequencing and Analysis

RNA from the lungs of control and experimental mice were sent for bulk sequencing separately. Approximately 0.4 ug of total RNA was used for library preparation. mRNA enrichment and library preparation was performed using the Polyadenylated (PolyA+) mRNA Magnetic Isolation Module (New England Biolabs) and NEBNext Ultra II RNA Library Prep Kit for Illumina (New England Biolabs), following the manufacturer’s protocol. All 12 samples were then pooled together and sequenced in one lane using Illumina Novaseq 6000 platform with paired-end 150bp (table S1). The raw fastq files from RNA-seq were mapped to GRCm38 mouse genome reference using STAR (version 2.5) with default parameters. More than 90% (table S1) of sequenced paired-end reads (above 50M reads for each library) were mapped to the mouse genome by STAR(Dobin et al., 2013). HTSeq(Anders et al., 2015) (version 0.6.0) was used to quantify gene expression, with Ensembl GRCm38.96 as a reference. DESeq2 (version 1.24.0) was used to perform the differential expression analysis on the HTSeq quantified count per gene. The top 200 up and down regulated genes (table S2), ranked by P-value from low to high with fold-change above 1.5 (or log2(fold-change) > 0.58), were used for biological process enrichment analysis in the DAVID database. Biological process terms with at least 5 differentially expressed genes and BHQ < 0.15 were selected. For aggregating redundant biological processes, GOSemSim(Yu et al., 2010) (version 2.10.0) was used to calculate the semantic similarity (‘Jiang’ method from GOSemSim) between significant biological processes. Redundant biological processes were manually merged into biological process categories (table S3).

### Dissociation Protocol for Single Cell Sequencing

For scRNA-seq, a total of 3 biological replicates (3 IH and 3 controls in the first experiment and 2 of each in the other replicates). On the last day of exposure, the mice were sacrificed, lung harvested, and tissue immediately placed in ice-cold PBS. Dissociation of the pooled whole mouse lung for each group (IH vs RA) was performed as previously described(Guo et al., 2019). Briefly, minced lung was placed in collagenase/elastase/dispase digestion buffer (Sigma-Aldrick, St. Louis, MO; Worthington Biochemical, Lakewood, NJ). After mixing on ice for approximately 3 minutes, the lung was minced again. After resting the suspension, the supernatant was passed through a 30 μM filter. A *Bacillus licheniformis* mix (Sigma-Aldrich) was added to the cell suspension, mixed on ice for approximately 10 min, and passed through a 30 μM filter. The suspension was spun at 500 g for 5 min at 4°C. The pellet was rinsed with a red blood cell lysis buffer. This was again passed through a 30 μM filter and then spun at 500 g for 5 min at 4°C. The cell suspension was resuspended in PBS/BSA and manually counted with a hemocytometer. The volume was adjusted to obtain a final concentration of approximately 1000 cells/μL to be loaded to the 10X Chromium platform.

### scRNA-seq Library Construction and Sequencing

The single cell suspension was applied to the 10X Genomics Chromium platform (San Francisco, CA) to capture and barcode cells, as described in the manufacturer’s protocol. Libraries were constructed using the Single Cell 3’ Reagent Kit (v2 Chemistry). The completed libraries were then sequenced using HiSeq 2500 (Illumina, San Diego, CA) running in Rapid Mode. Each sample was loaded onto two lanes of a Rapid v2 flow cell.

### scRNA-seq Data Processing

Raw data from 10X Genomics were demultiplexed and converted to a fastq file using cellRanger (v2.1.1) mkfastq. Reads from the same library sequenced in different flow cells (technical replicates) were combined and aligned to the mm10 genome reference using cellRanger count. Summary sentences for statistical mapping are presented in table S4. The gene expression profiles for cells from the three biological replicates of the IH group were combined with cellRanger aggr and were run an unsupervised analysis using the software Iterative Clustering and Guide-gene Selection (ICGS) versions 2 (AltAnalyze version 2.1.2) to generate reference clusters using the program defaults with euclidean clustering(DePasquale et al., 2019). ICGS2 grouped 12,324 cells into 25 reference clusters based on the expression profiles of 1,480 selected marker genes (table S5). All cells from control and IH groups were then aligned to these 25 reference clusters using cellHarmony(DePasquale et al., 2019). Uniform Manifold Approximation and Projection (UMAP) calculation was run using integrated function in AltAnalyze -v2.1.2 with default parameters. For annotating the 25 reference clusters into known lung cell-types, we prepared a comprehensive marker gene list for known lung cell types. The sources of this marker gene list included information from the Mouse Cell Atlas, ToppGene, and Lung Gene Expression Analysis (LGEA)(Chen et al., 2009; Consortium et al., 2018; Du et al., 2017). Additionally, we manually collected cell marker genes from published scRNA-seq studies performed in mouse or human lung(Guo et al., 2019; Zilionis et al., 2019). One-tailed Fisher exact test was used to perform enrichment analysis between marker genes for each cluster and the curated reference markers of known lung cell types. Each cluster was manually assigned to a specific cell type based on the known cell type with the lowest BH(Benjamini & Hochberg, 1995) adjusted P-value (GO-Elite software)(Zambon et al., 2012). Those clusters corresponding to the same annotated cell type were manually joined as one cell type for downstream analyses (e.g. endothelial corresponding to four clusters). This process reduced the 25 reference clusters into 19 cell types (table S6). For testing the cell type composition difference of mouse lung between experimental and control groups, a centered log ratio transformation was performed on the percentage of each cell type before applying the t-test (two-tailed). The statistical P-value from the t-test was adjusted with the BH method.

### Pseudo-bulk RNA-seq Differential Expression Analysis

To identify differentially expressed genes in each lung cell type between the control and IH groups with multiple biologic replicates, all cells assigned to the same cell type were aggregated into a “pseudo-bulk” data library by library. For each library, the sum of the reads per gene from cells assigned to the same cell type were used to represent the cell-type-specific gene expression profiles. Percentage of cells expressed per gene were calculated as a fraction of cells with >=1 read(s) for the gene in each cell-type. Count per one million UMI (CPM) for each cell type was calculated as the (sum of reads per gene / sum of reads) * 1000000 for each library. Differential expression analysis was performed with DESeq2 for each cell-type, using the sum of reads per gene as input, with three replicates in each of the control and IH groups. Ranked by P-value from low to high, the top 200 up and down regulated genes (table S7) with fold-change above 1.2 (or log2(fold-change) > 0.26) were used for biological process enrichment analysis in the DAVID database. Selecting and aggregating biological processes (table S8) were performed as described above in the Methods section labeled, “*Bulk RNA Sequencing and Analysis*.” We further extracted those genes enriched in circadian rhythm and immune response, and selected well-established genes in each biological process to demonstrate their expression variation under IH exposure in each cell type based on literature searches.

### Endothelial Subpopulations Analysis

To improve the accuracy for classifying endothelial cells, we reran the AltAnalyze-ICGS2 clustering algorithm using only the 5,579 cells annotated to pulmonary vascular endothelial cells from IH exposure groups, followed by a realignment of all vascular endothelial cells from both control and IH groups to these clusters using cellHarmony. AltAnalyze-ICGS2 produced 6 clusters with 314 marker genes (table S9). We matched the ICGS2 selected marker genes with vascular endothelial subpopulation marker genes presented in a study from Travaglini et al(Travaglini et al., 2019). This annotated the clusters into 4 subpopulations of endothelial artery, vein, capillary aerocytes, and general capillary cells (table S10). We further aggregated all cells of the same vascular endothelial subpopulations into “pseudo-bulk” data library by library. DESeq2 was used to detect differentially expressed genes for each subpopulation between the control and IH exposure groups. To select differentially expression genes in each endothelial subpopulation, the cut-off was set as BHQ < 0.2 and fold-change > 1.2 (or log2(fold-change) > 0.26). Those differential expression genes (table S11) in endothelial capillary cells were used for BP enrichment analysis in the DAVID database. Selecting and aggregating biological processes (table S12) were performed as described above in the Methods section labeled, “*Bulk RNA Sequencing and Analysis*.”

### Association Analysis on IH Responsive Genes with Pulmonary Disease Genes and Drug Targets at the Cell-type Level

Gene-disease association information was downloaded from the DisGeNET(Piñero et al., 2017) database (curated gene-disease associations). The downloaded file was filtered with keywords to specifically select genes linked to pulmonary diseases, which includes“allergic asthma”, “asthma”, “chronic obstructive airway disease”, “pulmonary hypertension”, “chronic thromboembolic pulmonary hypertension”, “idiopathic pulmonary arterial hypertension”, “familial primary pulmonary hypertension”, “idiopathic pulmonary hypertension”, “interstitial lung fibrosis”, “pulmonary fibrosis”, and “idiopathic pulmonary fibrosis”. One-tailed Fisher exact test was used to determine if the top 200 up or down regulated genes in each cell-type exposed to IH significantly overlapped with these pulmonary disease associated genes (table S13). The significant cutoff was set with a BH adjusted P-value from a Fisher exact test < 0.05 and at least 5 overlapped genes with any pulmonary disease gene set. For the association analysis between IH responsive genes and pulmonary drug targets, the xml file was downloaded from the DrugBank(Wishart et al., 2008) (version 5.1.4). The drugbankR package (https://github.com/yduan004/drugbankR) was used to parse the xml file to get each drug and its target genes. The parsed drug table was linked with the top 200 up or down regulated genes in each cell-type exposed to IH by drug target genes. The drug table was further filtered with respiratory tissue and disease-associated key words (e.g. asthma, lung, bronchus, airway et al.) to keep candidate drugs used to treat pulmonary diseases. The filtered table was manually curated to select drugs mainly indicated to treat pulmonary diseases. In the association analysis, the ‘homologene’ package (https://github.com/oganm/homologene) was used to find the human-unique homolog of mouse genes. CellPhoneDB is used to predict ligand-receptor interactions between cell-types. The pulmonary diseases relevant drug targeted genes list (Fig 6C) is used to select ligand-receptor interaction pairs. Genes with unknown drug effects were filtered out from the analysis. Human homolog gene is used for this analysis. Gain of interactions and loss of interactions is calculated by summing up the number of ligand-receptor interactions that were only significantly present in either hypoxia or control (P-value < 0.05 and mean > 0.1), respectively, for each cell pairs comparison. Two-tailed Fisher exact test was used to calculate significance changes of each cell type. Hive plot is made using the HiveR package (https://github.com/bryanhanson/HiveR).

### Lung Fixation, Histological Staining, Immunofluorescence, and Confocal Microscopy

For histological staining, mice were sacrificed, and lung inflation fixation was immediately performed. After exposure of the trachea and lungs, the trachea was cannulated with a scalp vein cannula (EXELINT, Redondo Beach, CA), and 10% neutral buffered formalin was gravity-perfused into the lung at a height of 25 cm. After infusion, the lung was harvested and placed in formalin for 24 hours. Whole lung was then dehydrated in 70% ethanol and embedded in paraffin. For morphologic evaluation, 5 μm thick sections were cut from the paraffin blocks and stained with hematoxylin and eosin.

Immunofluorescence staining on 10% formalin fixed mouse lung was performed on 5 μm thick paraffin embedded tissue sections. Tissue slides were melted at 60°C for two hours, following rehydration through xylene and alcohol, and finally in PBS. Antigen retrieval was performed in 0.1 M citrate buffer (pH 6.0) by microwaving. Slides were blocked for 2 hours at room temperature using 4% normal donkey serum in PBS containing 0.2% Triton X-100, and then incubated with primary antibodies diluted in blocking buffer for approximately 16 hours at 4°C. Primary antibodies included ACTA2 (1:2000, Sigma-Aldrich), EMCN (1:200, R&D Systems), FOXFI (1:100, R&D Systems), HOPX (1:100, Santa Cruz Biotechnology), MKI67 (1:100, BDBiosciences), POSTN (1:100, ABCAM), Pro-SFTPC (1:1000, Seven Hills Bioreagents), SOX9 (1:100, Millipore), and LYVE1 (1:100, ABCAM). Secondary antibodies conjugated to Alexa Fluor 488, Alexa Fluor 568, or Alexa Fluor 633, were used at a dilution of 1:200 in blocking buffer for 1 hour at room temperature. Nuclei were counterstained with DAPI (1μg/ml) (Thermo-Fisher). Sections were mounted using ProLong Gold (Thermo-Fisher) mounting medium and coverslipped. Tissue sections were then imaged on an inverted Nikon A1R confocal microscope using a NA 1.27 objective using a 1.2 AU pinhole. Maximum intensity projections of multi-labeled Z stack images were generated using Nikon NIS-Elements software.

## Supporting information

DatafileS1

DatafileS2

DatafileS3

DatafileS4

## Acknowledgements

We thank K.A. Wikenheiser-Brokamp for her thoughtful discussion and guidance. We thank Bruce J. Aranow, Yan Xu, Minzhe Guo, Kashish Chetal, and Emily R. Miraldi for their thoughtful discussion about scRNA-seq analysis. We would like to thank S. Steven Potter for allowing us to use resources available in his lab. We would also like to thank Kalpana Srivastava for work on sectioning tissue specimens and completing immunofluorescence assays.

## Author contributions

G.W., Y.Y.L., E.M.G., M.D.R., L.J.F., J.B.H., and D.F.S. designed research; G.W., Y.Y.L., E.M.G., A.P., J.K., L.J.F., and D.F.S performed research; G.W., Y.Y.L., N.S., J.A.W., and J.B.H. contributed analytic tools; G.W., Y.Y.L., E.M.G., M.D.R., N.S., J.A.W., L.J.F., and D.F.S. analyzed data; G.W., Y.Y.L., M.D.R., L.J.F., J.B.H., and D.F.S. wrote this paper.

## Data and Materials Availability

All data associated with this study are present in the paper or the Supplementary Materials and have been incorporated into GEO database (GSE145436).

## Supplemental figures

**Fig. S1.**
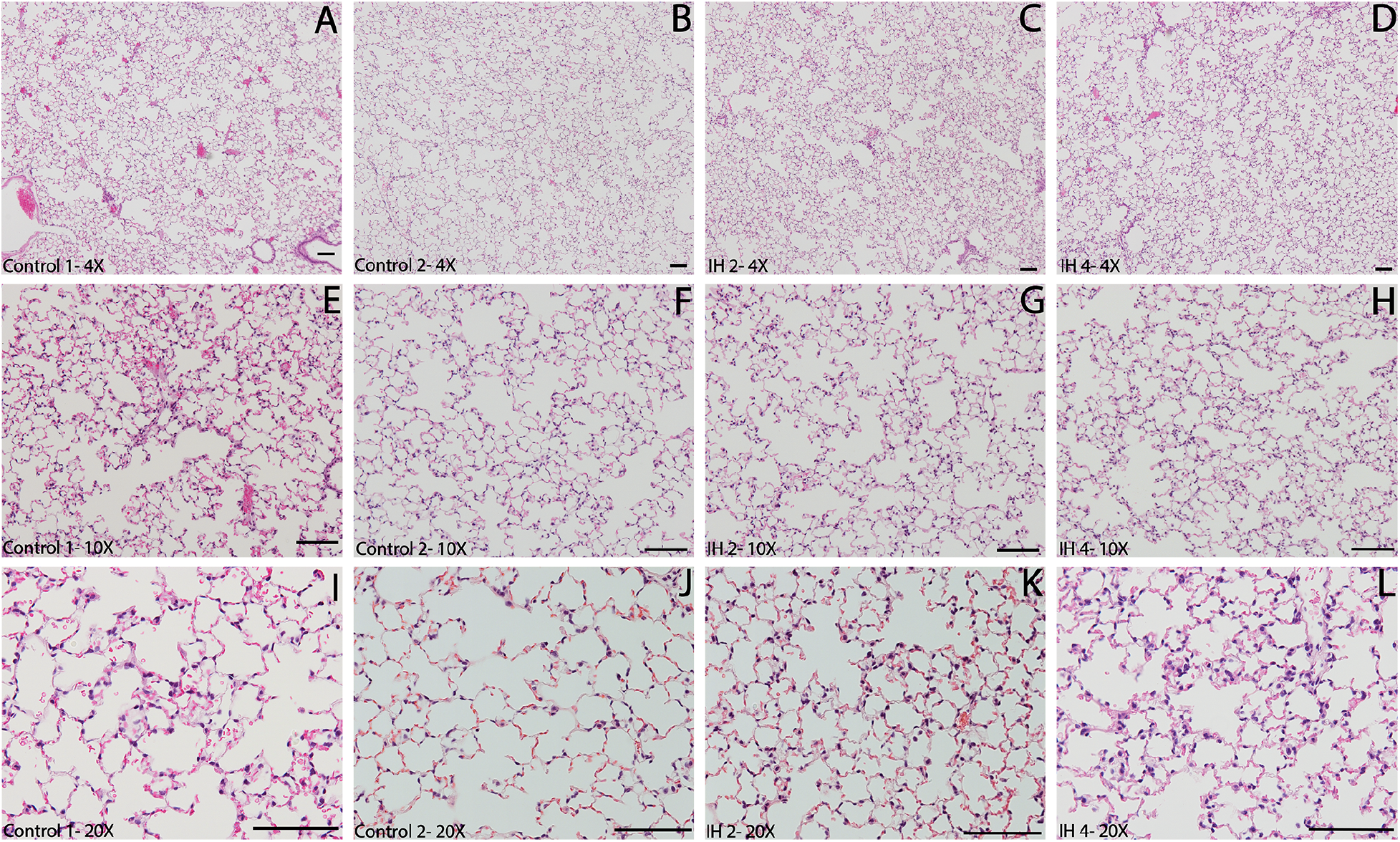
Hematoxylin and eosin–stained sections of whole lung from mice exposed to IH vs Controls do not reveal gross histologic changes. Two representative IH and two control mice with HE stained sections were shown at 4x (A-D), 10x (E-H) and 20x (I-L) magnification. The scale bars all correlate to 100 μm width.

**Fig. S2.**
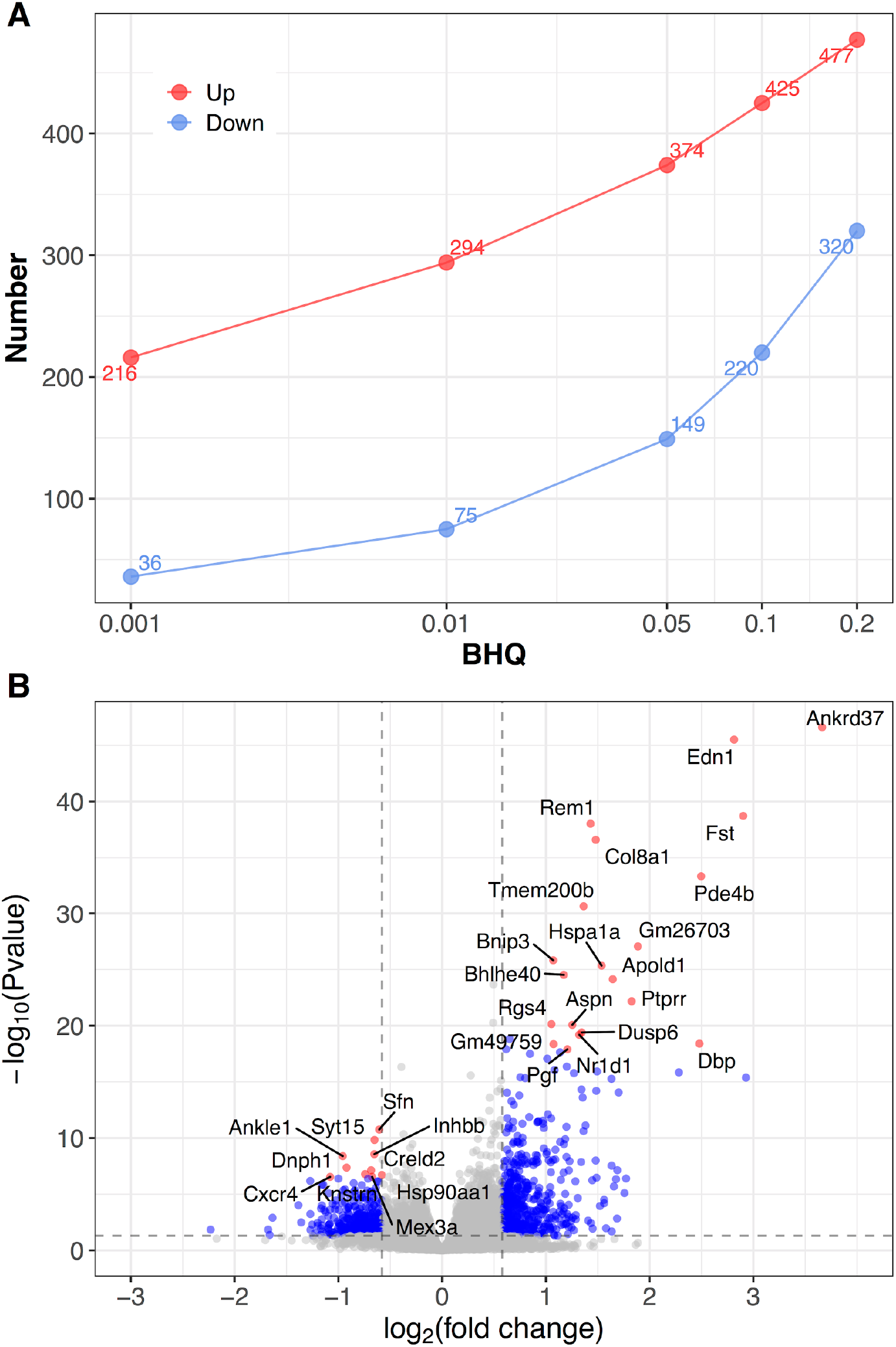
Differentially expressed genes from bulk RNA-seq analysis of whole lung from mice exposed to IH vs controls. (A) At series BHQ cut-offs, the number of up and down regulated genes in response to IH are shown with red and blue lines, respectively. (B) A volcano plot shows the differentially expressed genes from lung of mice exposed to IH. Ranked by P-value, top 20 up regulated genes and top 10 down regulated genes were labeled.

**Fig. S3.**
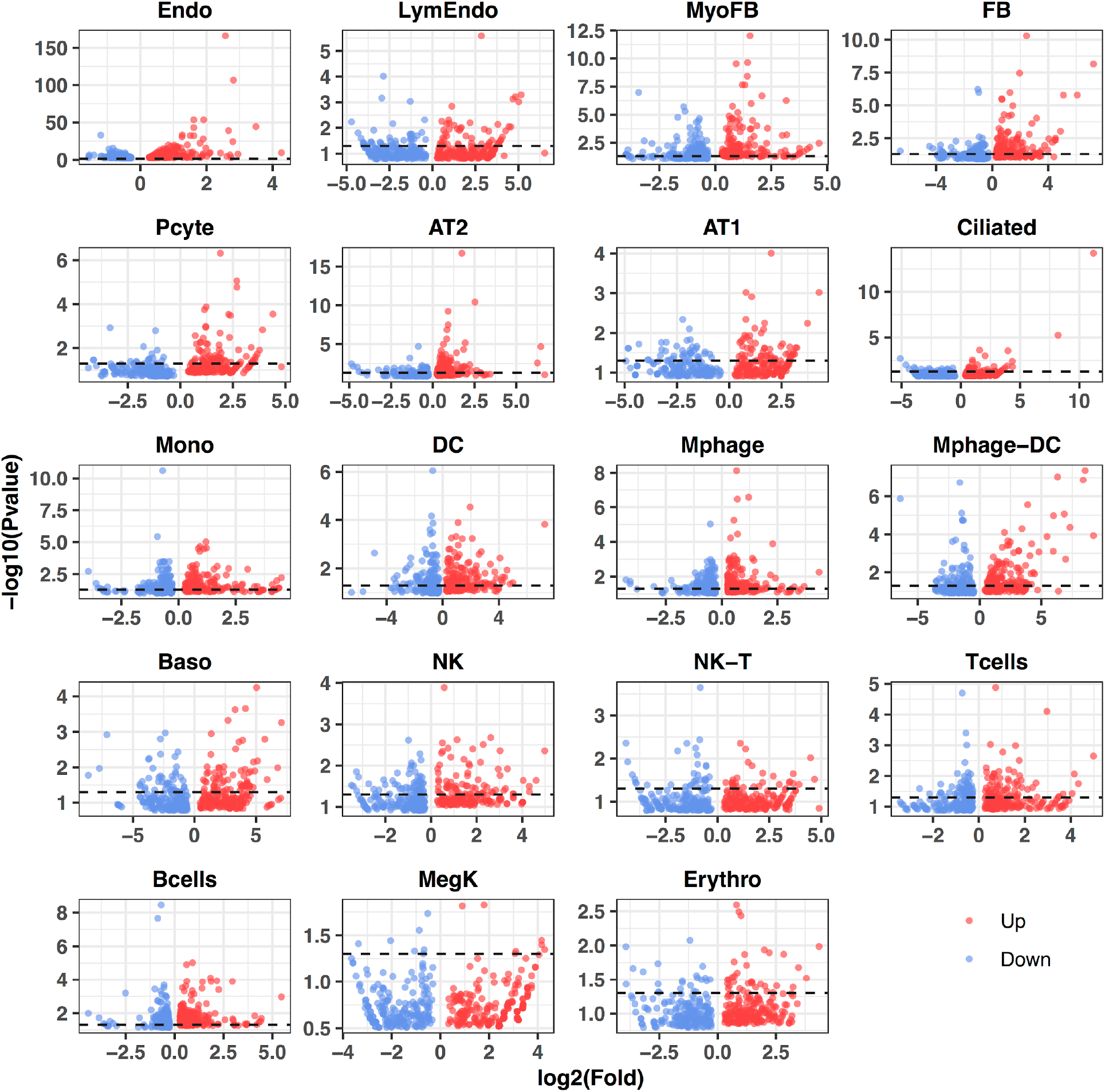
P-value distribution of the top 200 up and down regulated IH-responsive genes in 19 lung cell types. Each red or blue point indicates one up or down regulated gene. The black dashed line indicates a P-value of 0.05.

**Fig. S4.**
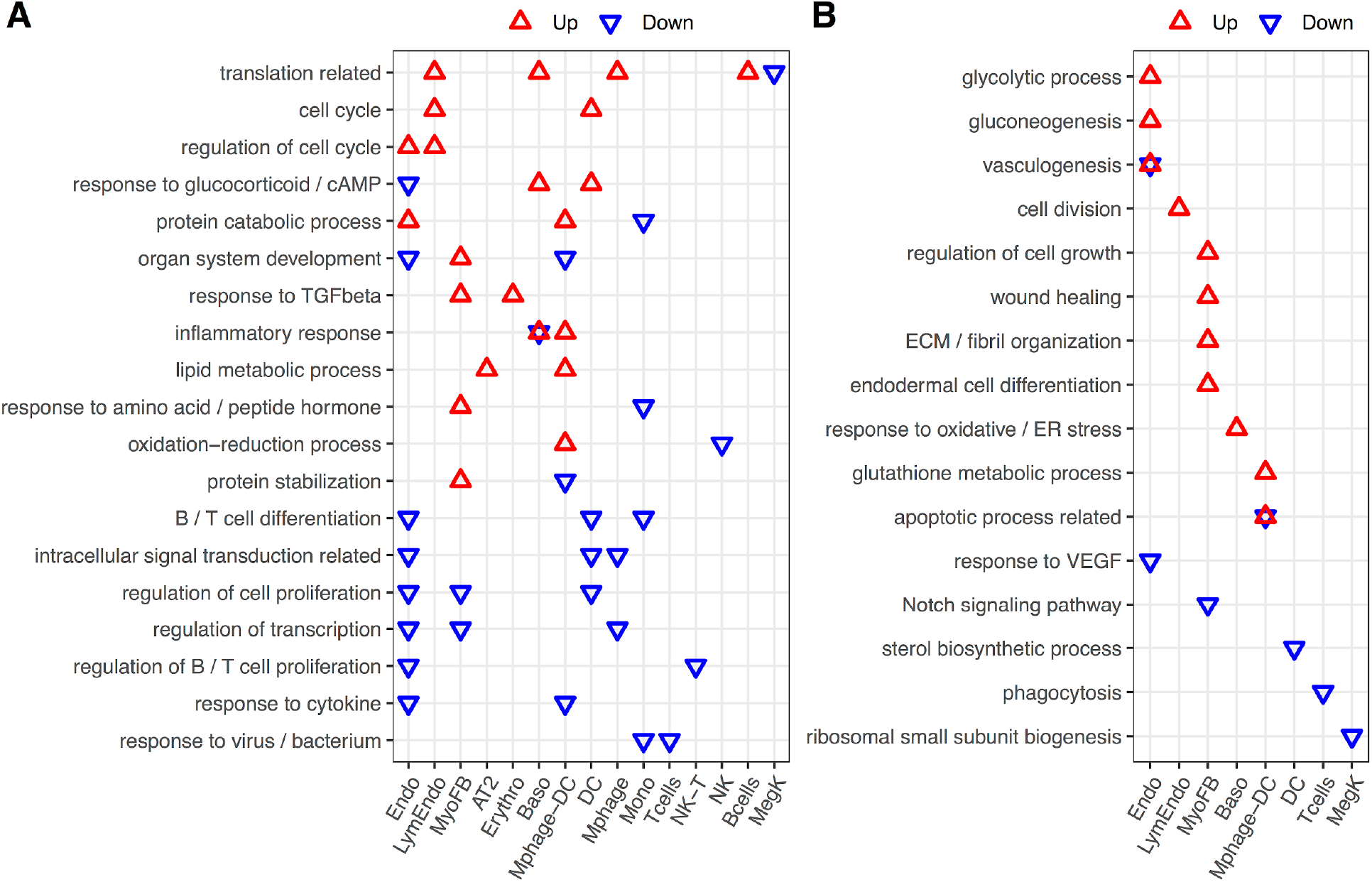
Enriched biological processes from differentially expressed genes from whole mouse lungs exposed to IH vs controls. Enrichment analysis was performed in the DAVID database using the top 200 up and down regulated genes identified from each cell type. Redundant biological processes are merged into one category. Biological processes enriched in up and down regulated genes are indicated by red and blue triangles, respectively.

**Fig. S5.**
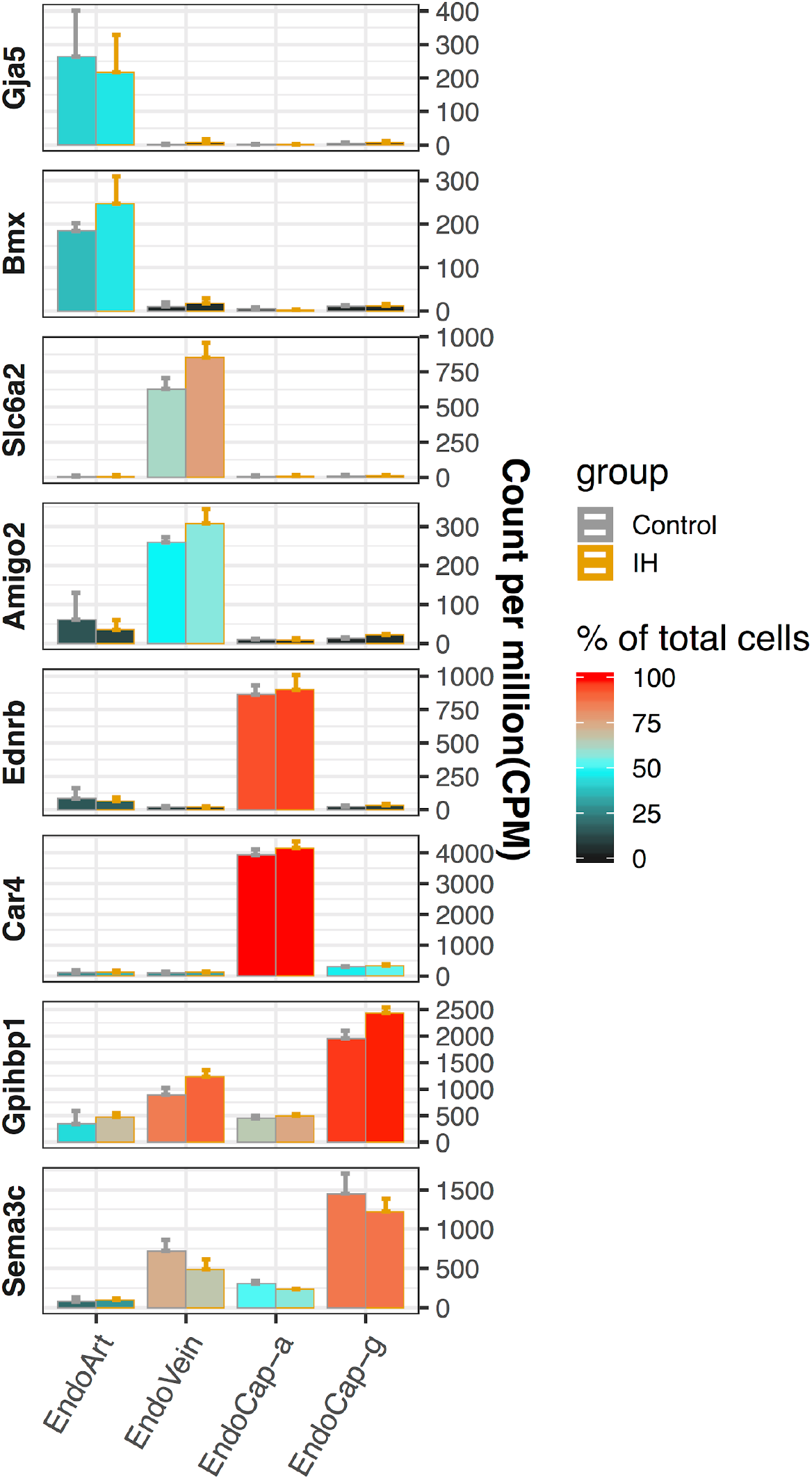
Validation of endothelial subpopulation assignments with known markers. The marker genes collected from the literature and assigned endothelial subpopulations are listed on the y-axis and x-axis, respectively. The average CPM (count per million reads) of three replicates from control (grey) and IH (orange) mice are indicated by the height of the bar. The error bars indicate standard deviation based on the three biological replicates. The average percentage of cells with detected expression of each marker gene among the endothelial subpopulations is indicated by the heatmap color. The four endothelial subpopulations include endothelial artery (EndoArt), vein (EndoVein), capillary aerocytes (EndoCap-a), and capillary general (EndoCap-g).

**Fig. S6.**
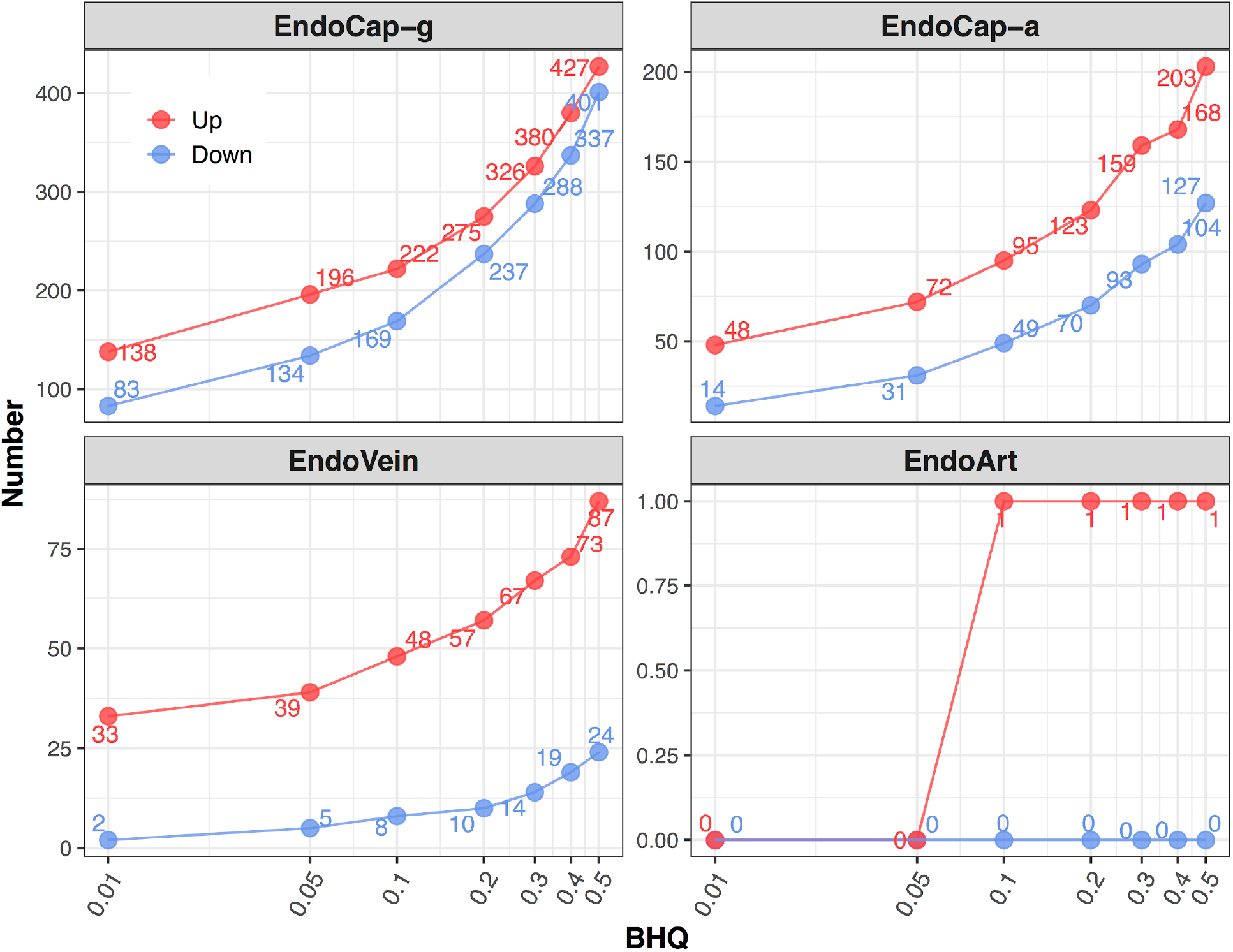
Number of differentially expressed IH-responsive genes in four lung endothelial subpopulations using BHQ cut-offs. At series BHQ cut-offs, the number of up and down regulated genes in lung endothelial subpopulations as a response to IH are shown with red and blue lines, respectively. The four endothelial subpopulations include endothelial artery (EndoArt), vein (EndoVein), capillary aerocytes (EndoCap-a), and capillary general (EndoCap-g).

**Fig. S7.**
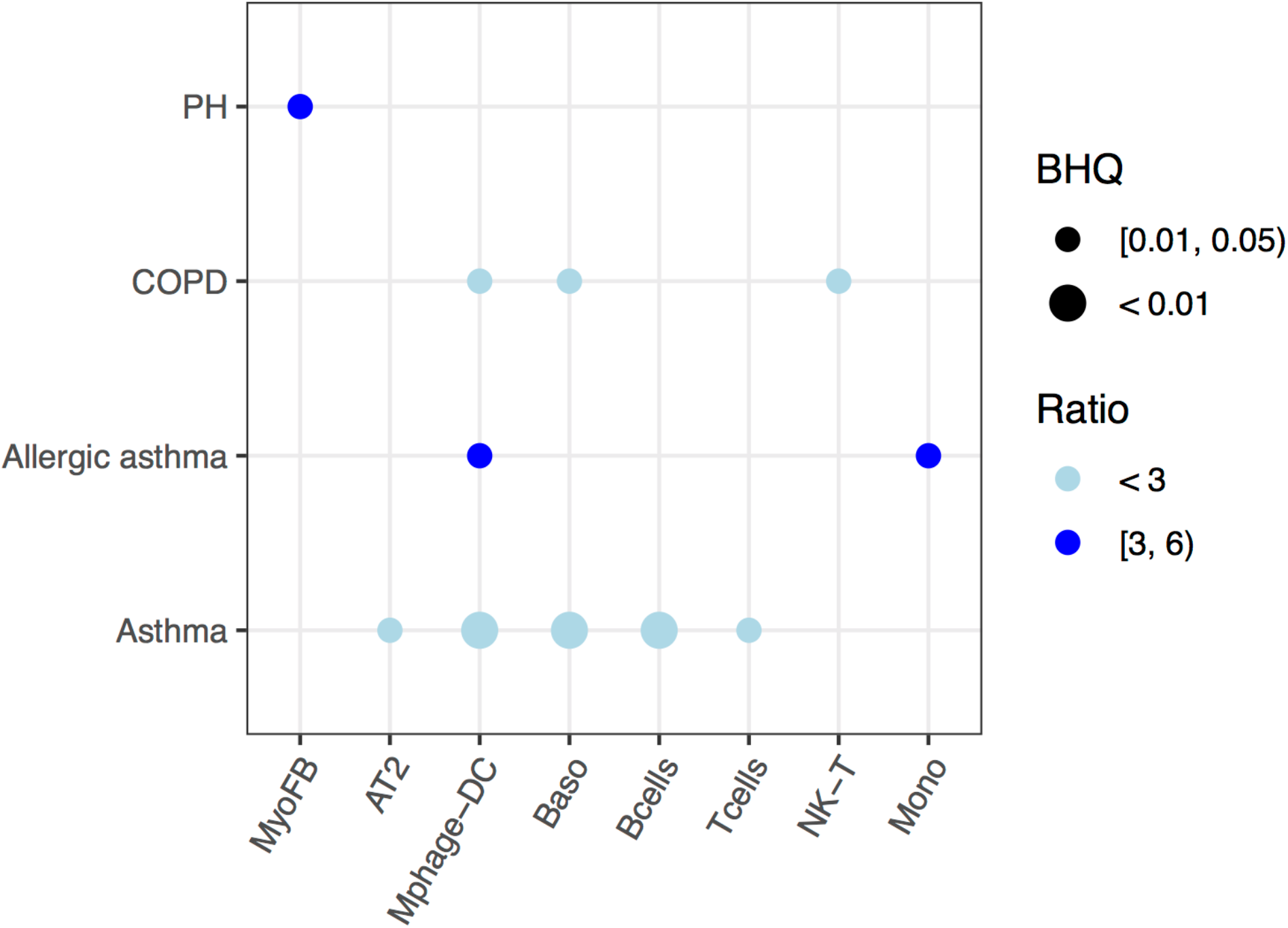
Pulmonary disease-associated genes are enriched in down regulated genes in different cell types in the lung from mice exposed to IH. Enrichment analysis was performed in the DisGenet database, using the top 200 down regulated genes identified from each cell type. The average fold change of down regulated genes in each cell type for the same disease is indicated by color scale. The point size indicates the enrichment BHQ value from the fisher exact test. The point color indicates the enrichment ratio. The pulmonary diseases include asthma, allergic asthma, chronic obstructive airway disease (COPD), and pulmonary hypertension.

**Fig. S8.**
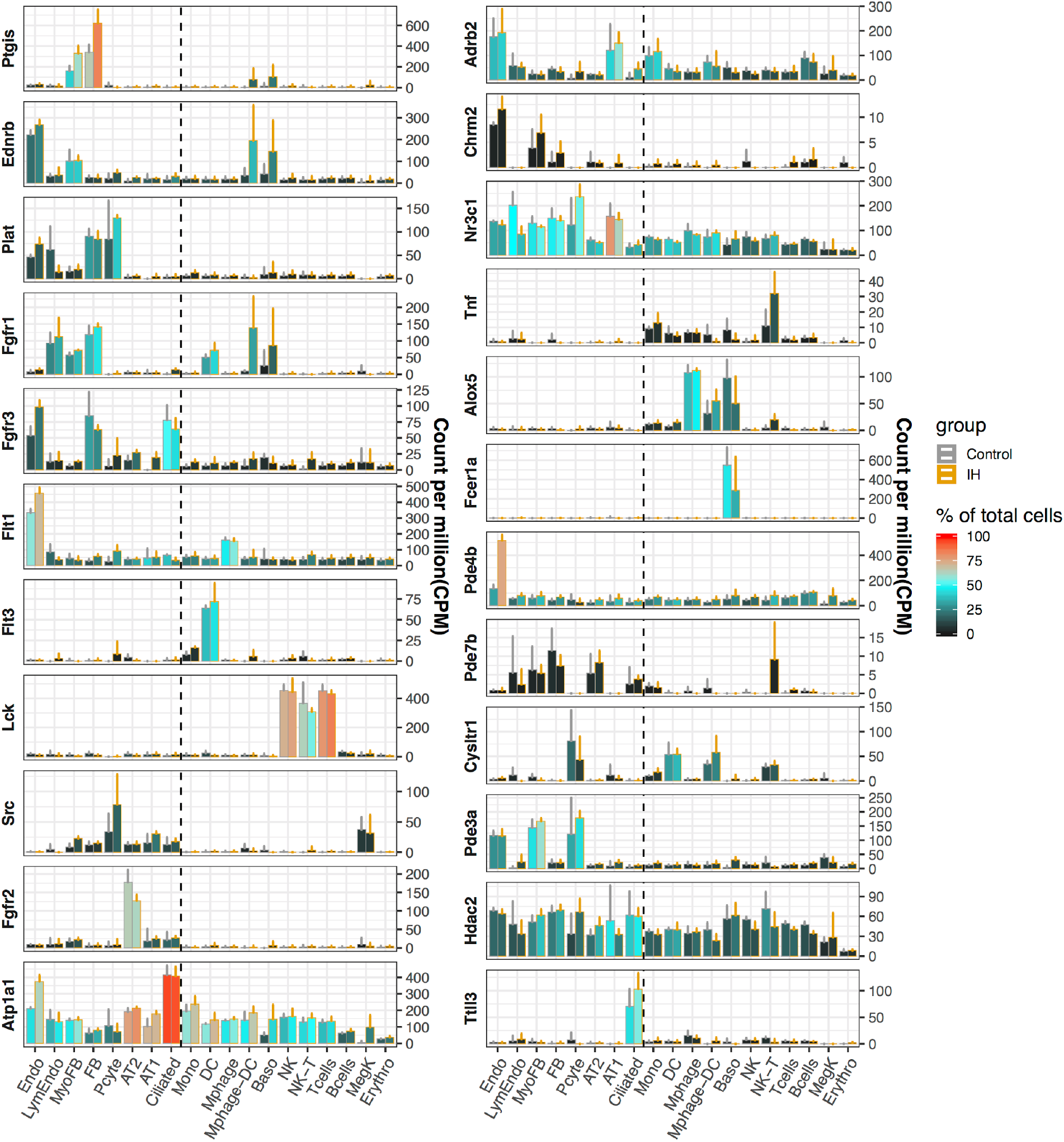
Pulmonary drug targets with cell type specific expression profiles in mouse lung. The average CPM (count per million reads) of three control (grey) and IH (orange) replicates are indicated by the height of the bar. The error bar indicates standard deviation from the three biological replicates. The average percentage of cells with detected expression of each marker gene among specific cell types is indicated by the color scale. The list of cell types include: endothelial cells (Endo), B cells (Bcells), natural killer cells (NK), T cells (Tcells), natural killer cells (NK), natural killer T cells (NK-T), macrophages (Mphage), basophils (Baso), monocytes (Mono), macrophages-dendritic CD163+ cells (Mphage-DC), dendritic cells (DC), megakaryocytes (MegK), fibroblasts (FB), myofibroblasts (MyoFB), pericytes (Pcyte), alveolar type I cells (AT1), lymphatic endothelial cells (LympEndo), erythroblasts (Erythro), alveolar type II cells (AT2), and ciliated cells (Ciliated).

**Fig. S9.**
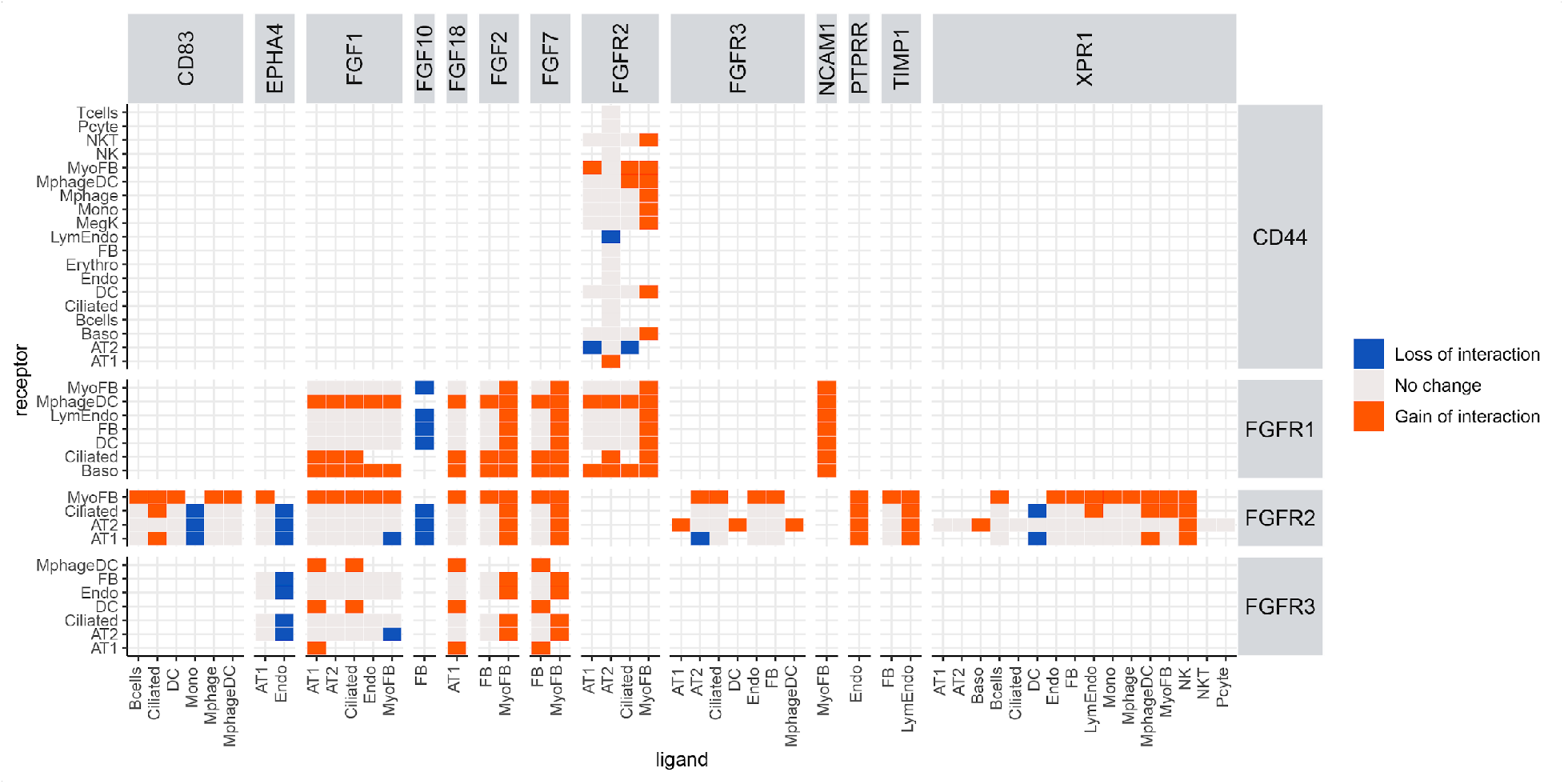
Ligand-receptor interaction analysis reveals a major role for myofibroblasts in activating the FGF signaling pathway as a response to IH. Heatmap shows ligand-receptor pair interaction changes in different cell pairs. Orange indicates a new interaction found in response to IH but not in control samples (gain of interaction), and blue indicates an interaction found in controls but not in response to IH (loss of interaction). Myofibroblasts show a gain of interaction between FGF2, FGF7 and NCAM1 to the FGF receptor in multiple cell types, indicating its role in activating the FGF signaling pathway in response to IH.

**Other Supplementary Material for this manuscript includes the following:**

**Data file S1 contains the following supplementary tables:**

Table S1 (Microsoft Excel format). The alignment statistics of bulk RNA-seq data for IH exposure and control mice.

Table S2 (Microsoft Excel format). Top 200 up and down regulated genes in mice lung under IH exposure from the bulk RNA-seq data.

Table S3 (Microsoft Excel format). DAVID enriched biological processes and merged categories using top 200 differential expression genes from the bulk RNA-seq data.

**Data file S2 contains the following supplementary tables:**

Table S4 (Microsoft Excel format). The alignment statistics of scRNA-seq data for IH exposure and control mice.

Table S5 (Microsoft Excel format). The marker genes list for 25 clusters from AltAnalyze analysis of scRNA-seq data from IH exposure mice.

Table S6 (Microsoft Excel format). The annotated cell types for 25 cellHarmony aligned clusters of scRNA-seq data from IH exposure and control mice.

Table S7 (Microsoft Excel format). Top 200 up and down regulated genes in 19 detected lung cell types under IH exposure from the scRNA-seq data.

Table S8 (Microsoft Excel format). DAVID enriched biological processes and merged categories using top 200 differential expression genes in each lung cell type under IH exposure.

**Data file S3 contains the following supplementary tables:**

Table S9 (Microsoft Excel format). The marker genes list for 6 clusters from AltAnalyze analysis of scRNAseq-data from annotated vascular endothelial cells of IH exposure mice.

Table S10 (Microsoft Excel format). The annotated endothelial subpopulations for 6 cellHarmony aligned clusters of scRNA-seq data from annotated vascular endothelial cells of IH exposure and control mice.

Table S11 (Microsoft Excel format). Differential expression genes of 4 annotated endothelial subpopulations under IH exposure from the scRNA-seq data.

Table S12 (Microsoft Excel format). DAVID enriched biological processes and merged categories using differential expression genes of lung capillary cells under IH exposure.

**Data file S4 contains the following supplementary tables:**

Table S13 (Microsoft Excel format). The pulmonary disease associated genes from top 200 up or down regulated genes in each detected lung cell-type exposed to IH.

